# Beyond parallel evolution: when several species colonize the same environmental gradient

**DOI:** 10.1101/662569

**Authors:** Alan Le Moan, Oscar Gaggiotti, Romina Henriques, Paulino Martinez, Dorte Bekkevold, Jakob Hemmer-Hansen

## Abstract

Genomic signatures associated with population divergence, speciation and the evolutionary mechanisms responsible for these are key research topics in evolutionary biology. Evolutionary radiations and parallel evolution have offered opportunities to study the role of the environment by providing replicates of ecologically driven speciation. Here, we apply an extension of the parallel evolution framework to study replicates of ecological speciation where multiple species went through a process of population divergence during the colonization of a common environmental gradient. We used the conditions offered by the North Sea – Baltic Sea environmental transition zone and found clear evidence of population structure linked to the Baltic Sea salinity gradient in four flatfish species. We found highly heterogeneous signatures of population divergence within and between species, and no evidence of parallel genomic architecture across species associated with the divergence. Analyses of demographic history suggest that Baltic Sea lineages are older than the age of the Baltic Sea itself. In most cases, divergence appears to involve reticulated demography through secondary contact, and our analyses revealed that genomic patterns of divergence were likely the result of a combination of effects from past isolation and subsequent adaptation to a new environment. In one case, we identified two large structural variants associated with the environmental gradient, where populations were inferred to have diverged in the presence of gene flow. Our results highlight the heterogeneous genomic effects associated with complex interplays of evolutionary forces, and stress the importance of genomic background for studies of parallel evolution.

## Introduction

Understanding the evolutionary history and the mechanisms involved in population divergence and speciation is a central research question in evolutionary biology. Genomic data have facilitated genome wide inferences of evolutionary forces acting on the process of divergence, and have provided unprecedented resolution to characterise complex interactions of demographic history and natural selection, as natural populations diverge and adapt to new environmental conditions. Here, we use a comparative framework to study divergence in marine fishes that have successfully colonized an extreme environment in recent evolutionary time. We show that signals of divergence are highly heterogeneous among species and likely result from the complex interplay of a demographic history that is older than the regions they now inhabit and more recent adaptation to the extreme environment.

While limited general inference can be drawn from single case studies, comparative frameworks offer powerful approaches for providing insights on the processes of population divergence and speciation (Burri, 2017; Cruickshank and Hahn, 2014; Galtier, 2019; Roux *et al.*, 2016). Such frameworks have been used to understand the effect of the environment when speciation occurs in the face of gene flow (Nosil, 2012; Rundle and Nosil, 2005). Specifically, the comparison of several speciation events linked to common environmental pressures allows us to characterize the repeatability of evolution through the identification of similar adaptive characteristics. These similarities correspond to convergent evolution if they evolved independently in two or more replicates of ecologically driven speciation (Losos, 2011).

Two classical frameworks have been used to study replicated ecological speciation in the face of gene flow (Elmer and Meyer, 2011). The first focuses on parallel evolution when a species colonizes and adapts to similar environmental contrasts across different geographically isolated areas (Johannesson, 2001; Schluter and Nagel, 1995). The second concerns ecological radiation following the colonization of underutilized niches by a single species (Seehausen, 2004). These frameworks have identified several cases of evolutionary convergence underlined by similar genetic pathways across a suite of organisms (Hench *et al.*, 2019; Hohenlohe *et al.*, 2010; Lamichhaney *et al.*, 2015; Muschick *et al.*, 2012; Ravinet *et al.*, 2016). The inferences from genome-wide variation has recently highlighted the role of gene flow between recently diverged species as a fuel for evolutionary radiations (Malinsky *et al.*, 2018; Meier *et al.*, 2017), as well as selection on standing variation to promote the parallel evolution of ecotypes (Belleghem *et al.*, 2018; Marques *et al.*, 2019). Therefore, identifying a relevant framework with replicates of ecological divergence with independent genetic backgrounds (i.e. isolated species with complete allelic sorting) is still essential to disentangle the effects of selection in response to the environment from other evolutionary mechanisms (Foote, 2018; Lee and Coop, 2017). The repeated colonization of the same environmental gradient by several species thus provides a third approach to study independent replicates of ecological divergence.

Past events of climatic cycling can offer such opportunities by restoring new habitats that were previously inaccessible (Hewitt, 2000). The Baltic Sea basin represents an example of such area. It was entirely covered by ice during the Last Glacial Maximum (LGM), 55 000 years ago (Houmark-Nielsen and Kjær, 2003). The connection to the Atlantic marine environment was established approximately 8 000 years ago following glacial retreat (Björck, 1995). Important ecological gradients are found along the North Sea – Baltic Sea Transition Zone (NBTZ). For example, the North Sea is a fully marine environment, with salinity at 35 PSU, but the salinity decreases along the NBTZ leading to a gradient of increasingly brackish water, with moderate salinity (ca. 13 PSU) in the south-west of the Baltic Sea, and low salinity (ca. 3 PSU) in its northern parts (Janssen *et al.*, 1999). Several marine species have successfully colonized and adapted to this gradient that represents an extreme environment for many marine organisms, with several species showing clinal patterns of genetic differentiation along the NBTZ (Johannesson and Andre; Hemmer-Hansen *et al.*, 2007; Nielsen *et al.*, 2009). Comparative population genetic analyses have been performed in the area with the use of few genetic markers (Johannesson and André, 2006), but so far the comparative framework has not been extended with the use of higher genome coverage.

Here, we used a comparative population genomic approach to study the evolutionary processes involved during the colonization of the Baltic Sea. We studied population structure of four closely related (but genetically isolated) flatfish species: turbot *(Scophthalmus maximus),* common dab *(Limanda limanda),* European plaice *(Pleuronectes platessa)* and European flounder *(Platichthys flesus).* These species were considered replicates of population divergence and were analysed independently. Our analyses showed heterogeneous genetic patterns of North Sea (NS) – Baltic Sea (BS) differentiation across species, both for magnitudes and genomic patterns of differentiation. Most of the genetic differences found were strongly associated with the environmental gradient of the NBTZ. However, the majority of the genomic regions underlying the NS-BS divergence were not shared between species. The timing of divergence between NS-BS populations was inferred to be five to ten times older than the age of the Baltic Sea. In three species, population divergence was inferred to involve past isolation followed by a secondary contact phase, and our analyses revealed that highly diverged genomic regions were likely the result of a combination of past isolation and subsequent adaptation to the environmental gradient. The only exception was found in European plaice where sympatric divergence provided a better explanation for the observed pattern of differentiation. Interestingly, the plaice genomic outlier regions were clustered on two distinct linkage blocks, likely revealing the presence of structural variants (SVs) in the genome. Altogether, this study shows the heterogeneity of the evolutionary pathways involved during the colonization of an environmental gradient, and our results highlight the importance of the genomic background for population divergence and adaptation in these highly diverse species.

## Results and Discussion

### Population structure

In all four species, a clear genetic separation of the North Sea and Baltic Sea samples was observed (Figure 1, horizontal axes of all PCAs). Although most species were divided into two populations (Figure S1), one additional cluster was found for the flounder (Figure 1c, vertical axis; Figure S2) distinguishing two ecotypes with different spawning strategies (Solemdal, 1973): the pelagic spawner (PEL from sites 5 to 11) and the demersal spawner (DEM from site 12). Most of the genetic breaks between clusters were localized between samples within the NBTZ, except for the PEL-DEM ecotype of flounder that was located inside the Baltic Sea (Figure 2). Furthermore, except for the DEM flounder, the genetic breaks corresponded roughly to the location of the environmental transition zone (Figure 2 a, b), confirming results from earlier work in marine taxa in the region (Johannesson and André, 2006; Limborg *et al.*, 2009).

**Figure 1.**
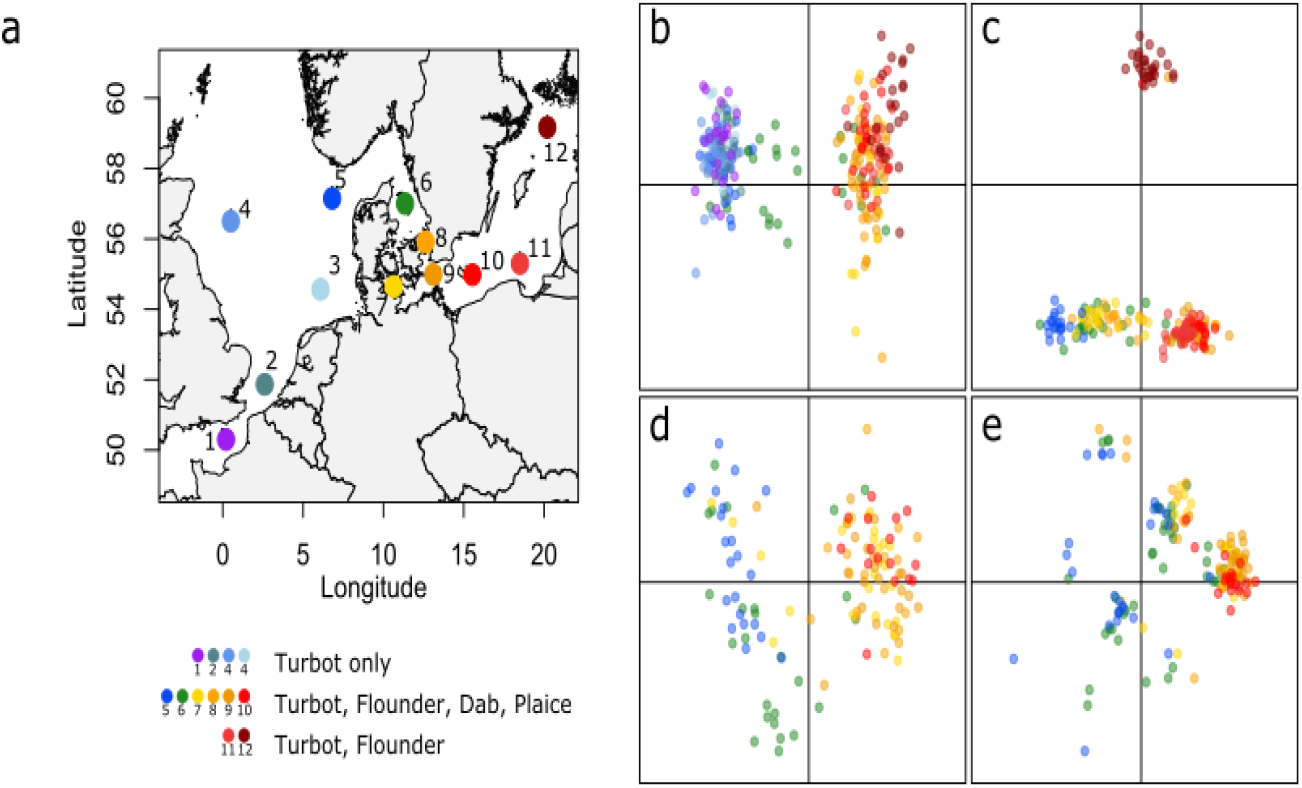
Geographical sampling of the four study species in a), and corresponding population structure in b)-e). Principal component analyses were performed separately for turbot (b), flounder (c), dab (d) and plaice (e). The colors of the individuals in b)-e) correspond to the colors of the sampling sites in a)

However, the extent of population admixture was variable across species which was supported by a general linear model (glm) analysis showing that the individual admixture proportions were significantly affected by the species, the geographical distance and the interaction between species and geographical distance (Chi^2^ = 19.44, p-value = 0.0006, df = 4, Figure 2a,b, Table S1). We did not find any evidence of hybridization between the two flounder ecotypes despite the presence of two DEM individuals sampled within the habitat range of the PEL ecotype (sites 9 and 10, Figure 1c and S3). The turbot showed limited NS-BS hybridization with only six admixed individuals, from site 6, assigned as F1 hybrids (Figure 1b and S3). In the remaining cases, the PEL flounder, the dab and the plaice displayed a continuum of hybridization along the NBTZ (Figure 1c-e, respectively) from which the admixed individuals could not accurately be classified as NS-BS hybrids (Figure S4). Consequently, the average NS-BS *F*_ST_ across all loci was two times higher in species showing limited hybridization (Figure S5, Flounder PEL-DEM *F*_ST_ = 0.044 and turbot *F*_ST_ = 0.039 tables S2 and S3) compared to species with substantial population admixture (Figure S5, flounder PEL-PEL *F*_ST_ = 0.013, plaice *F*_ST_ =0.014 and dab *F*_ST_ = 0.014, Table S2, S3, and S4).

**Figure 1:**
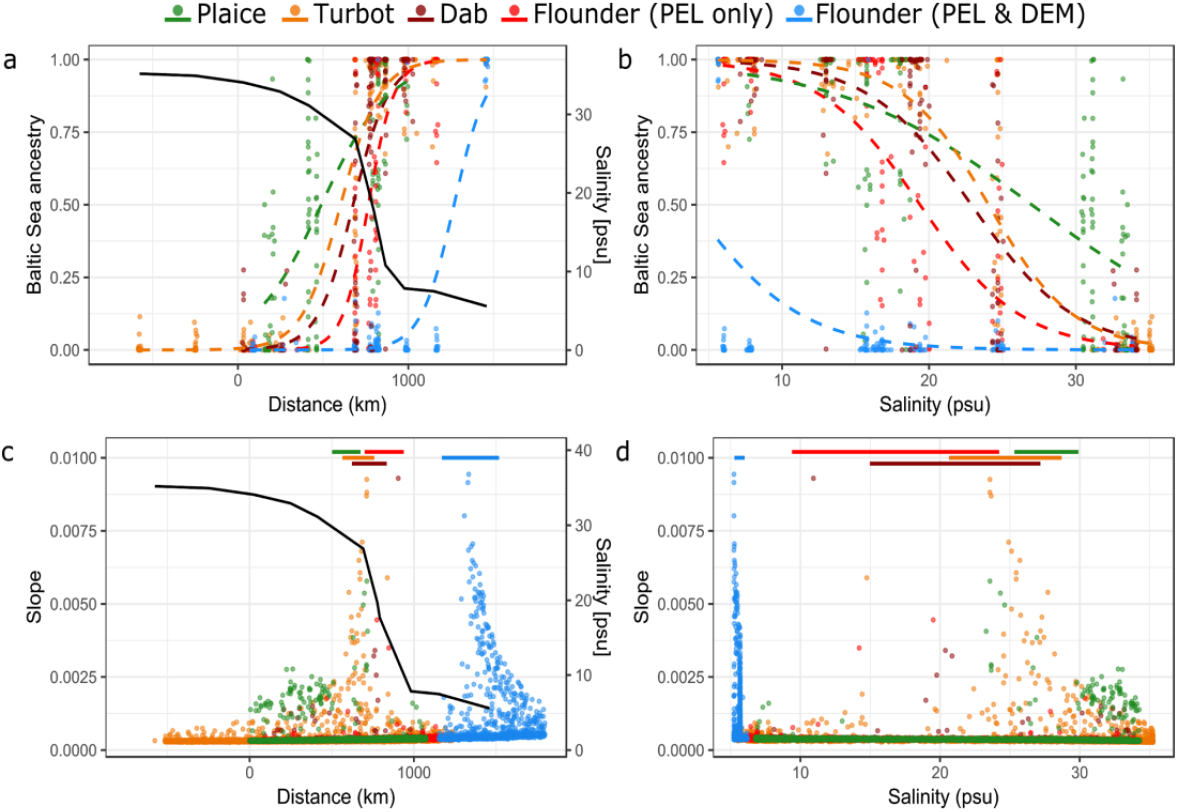
Geography of the genetic clines based on individual ancestry coefficients from cluster analyses (k=2) as a function of the sampling distance from the North Sea (a) and salinity (b); and the slope of allele frequencies for individual markers as a function of the distance of the cline center from the North Sea (c) and the salinity at the cline center (d). The dashed lines in a) and b) correspond to the fit of the general linear model. The solid black line in a) and c) shows the salinity along the North Sea Baltic Sea transition zone and the solid color lines (top of c) and d)) correspond to the confidence interval of the ancestry cline center estimated from data in a)

**Figure 2:**
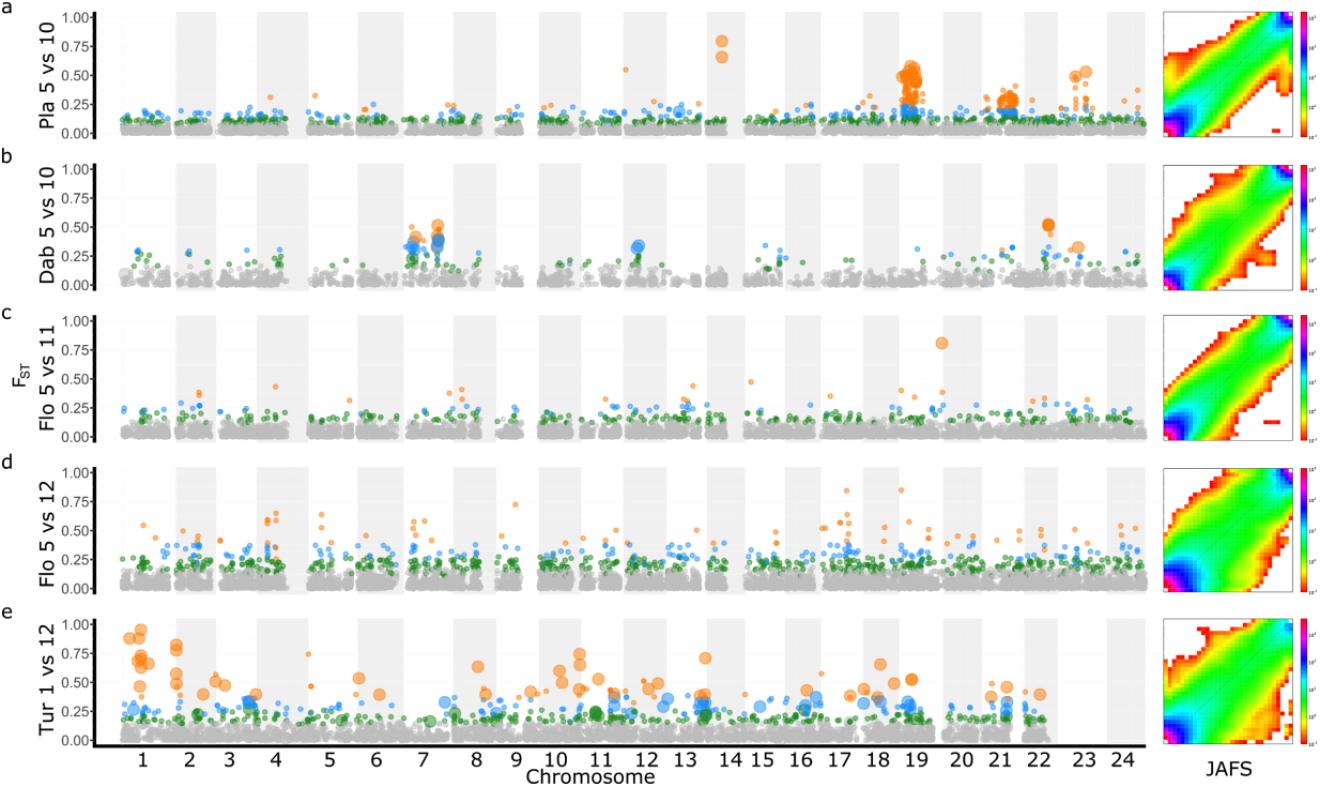
Manhattan plot of *F*_ST_ and observed Joint Allelic Frequency Spectrum (JAFS, with Baltic Sea on the x- and North Sea on the y-axis, respectively) between the two most geographically distant sampling sites for plaice (a), dab (b), pelagic flounder (c) pelagic vs. demersal flounder (d) and turbot (e). *F*_ST_ colors correspond to the 5% (green), 1% (blue) and 0.1% (orange) outliers from the *F*_ST_ selection test. Large dots represent *F*_ST_ outlier loci that were associated with salinity along the environmental gradient. No environmental association test was conducted for pelagic vs. demersal flounder

To further understand this inter-specific heterogeneity, we performed a genetic cline analysis across the environmental gradient to examine changes in allelic frequencies (estimated by steepness of slope and geographic location of slope center) along the environmental gradient. The variable extent of intra-specific differentiation reflected the number of loci associated with the structuring of the genetic clusters, more numerous and with steeper allelic clines between the PEL-DEM flounder and NS-BS turbot than between the other populations (Figure 2c,d). Altogether, these results suggest that NS-BS turbot and PEL-DEM flounder are more resistant to gene flow than the other NS-BS populations and/or that migration-drift equilibrium has not yet been reached in some of the populations in the other species. Higher genome wide differentiation for the turbot and PEL-DEM flounder was confirmed by the clear genome wide heterogeneity of *F*_ST_ in these comparisons (Figure 3d,e) compared to more localized peaks of *F*_ST_ across the genome in the remaining comparisons (Figure 3a-c)

Interestingly, only the plaice seems to show somewhat contrasting patterns across these analyses. Although this species showed low overall NS-BS differentiation, multiple highly differentiated markers with moderate value of allelic slope were detected (Figure 2c,d in green). These outliers were primarily clustered on two genomic islands of differentiation on chromosomes C19 and C21 (Figure 3a). High linkage disequilibrium (LD) was maintained over 8 Mbp along both islands (Figure S6), suggesting the presence of two large SVs with low recombination rates in the plaice genome. Interestingly, these SVs were already revealed by discrete groups on the PCA (Figure 1e), which corresponded to different combinations of the two alleles at the two putative SVs. Theoretically, they should result in a maximum of nine clusters (3 genotypes x 3 genotypes) from which only seven-eight were sampled in our study.

Sequences from the three Pleuronectidae species (plaice, dab and flounder) were aligned to the same genome, and therefore allowed us to explore the correlation between patterns of population diversity/divergence across species. The nucleotide diversity (π) and the net divergence *d*_XY_ (Cruickshank and Hahn, 2014) were weakly but significantly correlated in all the pairwise comparisons (Table S6), suggesting conserved recombination landscapes leading to similar variation of diversity along the genomes of the three species (Burri, 2017). However, we found no significant inter-specific correlation for sliding-widows average *F*_ST_ between North Sea and Baltic Sea lineages (Table S6), except between the flounder PEL and DEM lineages, which may reflect contemporary gene flow between the populations (Riquet *et al.*, 2019; Welch and Jiggins, 2014). Moreover, the 5% most differentiated regions (both with *F*_ST_ and *d*_XY_) were strongly and negatively correlated in all the species comparisons (Table S6). Thus, the regions that we detected to be highly differentiated were mostly private to each species (Figure 3a-d).

### Demographic inference

To understand the origin of the observed heterogeneous patterns of population structure across the four flatfish species, we compared observed data to 11 models of demographic history (Table S7 and S8) through analyses of the joint allelic frequency spectrum. In all cases, models with heterogeneous migration rates provided a better fit to the data, than models including neutral scenarios of divergence (Table 1). This suggests that the populations are experiencing a recent event of reproductive isolation resulting in heterogeneous gene flow along the genome (Wu, 2001), as also observed for the genome-wide distribution of loci with high *F*_ST_ (Figure 3).

**Table 1:** Optimal demographic scenarios inferred by diffusion approximation for the four study species. In order of appearance, the species, the first and second population used (Pop1 and Pop2 labelled as per Figure 1); the best demographic scenario (Model); the weighted AIC (wAIC) comparing the prediction of all the models and the parameters inferred: theta (θ), effective population size (N1 & N2); migration rate (Pop1 to Pop2 – m1>2 – and Pop2 to Pop1 – m2>1); reduced migration rate (mr); time since divergence between populations (T_S_); time of secondary contact (T_SC_), and the proportion (Q) of loci with reduced migration rates

| Species | Pop1 | Pop2 | Model | wAIC | $\theta$ | N1 | N2 | m1 | m2 | mr1 | mr2 | $T_s$ | $T_{sc}$ | Q |
| --- | --- | --- | --- | --- | --- | --- | --- | --- | --- | --- | --- | --- | --- | --- |
| Plaice | 5 | 10 | IM2m | 0.99 | 796 | 8.6 | 4.3 | 3.27 | 1.59 | 0.82 | 143.9 | 1.36 | - | 0.19 |
| Dab | 5 | 10 | SC2m | 1 | 657 | 19 | 0.69 | 6.61 | 0 | 0.2 | 26.8 | 2.01 | 0.44 | 0.25 |
| Flounder (PEL) | 5 | 11 | SC2m | 1 | 799 | 11 | 0.44 | 6.24 | 0.06 | 0.43 | 104.7 | 1.44 | 0.46 | 0.28 |
| Flounder (PEL-DEM) | 5 | 12 | SC2m | 0.99 | 892 | 6.5 | 1.7 | 5.82 | 9.04 | 1.02 | 5.2 | 1.31 | 0.11 | 0.29 |
| Turbot | 1 | 12 | SC2m | 0.99 | 1392 | 5.5 | 1.4 | 10.19 | 11.58 | 2.03 | 2.97 | 0.92 | 0.08 | 0.87 |

Except for plaice, for which the sympatric speciation model (IM) provided the best fit of the data, all the other demographic inferences suggested that a scenario of past isolation followed by secondary contact (model SC) was the most likely model to describe the origin of the observed population divergence (Table 1, S7 and Figure S8). These different scenarios could help explain the shape of the ancestry clines, which is somewhat sigmoidal in the cases of secondary divergence while linear in the case of primary divergence, as observed in plaice (Figure 2a). Using a generation time of 3.5 years (Erlandsson *et al.*, 2017) and a mutation rate of 10^-8^ per generation (The 1000 Genomes Project Consortium, 2010), time since divergence in all species was inferred to date roughly to the beginning of the LGM, more than 50 000 years ago (Table S8). Time since the secondary contact was estimated to be more recent, (around 10% of the total divergence time at approximately 5 000 years ago) between the two populations of turbot and the PEL-DEM ecotypes of flounder than in the dab and the PEL flounder (>30% of the total divergence, at approximately 15 – 22 000 years ago) (Table S8). Long periods of isolation can promote the accumulation of loci involved in reproductive barriers, such as Bateson-Dobzhansky-Muller incompatibilities (Dobzhansky, 1970). The variation in the estimates of isolation time inferred across species could therefore explain the different permeability to gene flow observed between clusters. Altogether, these results highlight that the origin of the North Sea and the Baltic Sea populations in the study species may involve several glacial refugia, and that the Baltic Sea populations may have diverged before the establishment of the Baltic Sea itself (Sick, 1965). Cases of ecological divergence following a temporary isolation have also been described in both parallel evolution (Le Moan *et al.*, 2016; Rougemont *et al.*, 2017; Rougeux *et al*., 2017) and evolutionary radiation (Foote *et al.*, 2016; Grant and Grant, 2009; Martin *et al*., 2015) scenarios. Complex demographic history associated with the divergence of Baltic Sea populations has been suggested in previous studies in the area (Johannesson and André, 2006; Nielsen *et al.*, 2004; Riginos and Cunningham, 2005; Sick, 1965). However, to our knowledge, this study provides the first formal test of this hypothesis with genomic data in the context of divergence of marine fish of the Baltic Sea.

The divergence times presented here are ten times higher than the previous estimation inferred from similar genomic data on the three lineages of flounder (Momigliano *et al.*, 2017). Although different methods were used for demographic inferences (Approximate Bayesian Computation vs diffusion approximation), and different models were tested (three-populations vs. pairwise inferences), continuous gene flow and the scenario of secondary contact were not included in previous work, which may explain the different conclusions reported here for the demographic modelling. The existence of multiple glacial refugia represents one potential explanation for the occurrence of several lineages, and similar patterns of discrete population clustering have also been observed in Atlantic cod *(Gadus morhua),* herring *(Clupea harengus)* and sandeel *(Ammodytes tobianus),* which are found across the North Sea and the Baltic Sea region (Barth *et al.*, 2017; Fietz *et al.*, 2018; Limborg *et al.*, 2012).

### Evidence of selection

To investigate signals of selection in the data, two different approaches were used to account for possibly “noise” associated with demographic history processes. First, we performed an *F*_ST_ outlier genome scan (Figure S8) by using the “neutral” parameters of divergence estimated by the best demographic scenario, in order to simulate the neutral envelope of differentiation (Beaumont and Nichols, 1996). Second, we used the programme Bayenv to detect loci with variation in allele frequencies that showed stronger association with environmental parameters than the overall population structure (Günther and Coop, 2013). Although fewer outliers were detected with the environmental association than with the genome scan approach (Table 2), both analyses showed heterogeneous outlier patterns within and between species (Figure 3). In general, environmental outliers were among the top 1 % or 0.1 % of the genome scan outliers (Figure 3 and S8, large dots in blue and orange). These outliers generally also displayed high values of allelic slope (Figure S9 and Figure S10) and the center of the outlier allelic clines were generally found within the confidence interval of the ancestry cline estimated with the glm analyses (Figure 2b and Figure S11). These findings suggest that signatures of selection may result from local adaptation (Hemmer-Hansen *et al.*, 2007; Johannesson and André, 2006) and/or genetic incompatibilities, which are likely to evolve during the isolation phase in a secondary contact scenario, and which can subsequently be trapped by environmental gradients through a coupling effect with local adaptation (Barton, 1979; Bierne *et al.*, 2011). Hence, it may be difficult to disentangle clearly the underlying mechanism of selection in these species.

**Table 2:**
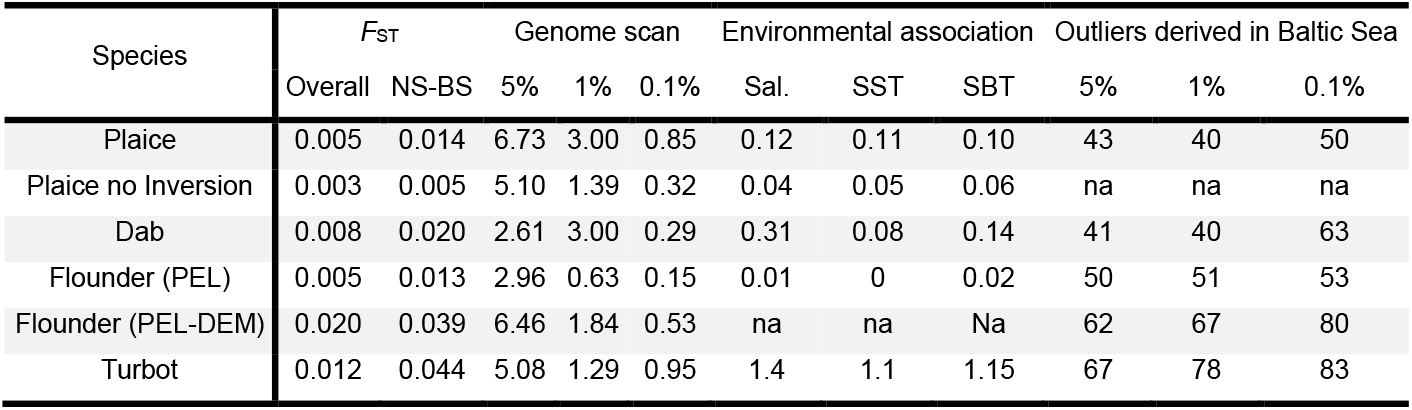
Summary statistics for population differentiation and the selection tests in the four species studied. In order of appearance, for each species, overall *F*_ST_, pairwise *F*_ST_ between the two most distant sites, the percentage of loci classified as outliers in the *F*_ST_ outlier scan for three different thresholds, the percentage of loci associated with 3 environmental factors (Salinity, Sea Surface Temperature and Sea Bottom Temperature), and the percentage of derived mutations within the Baltic Sea lineage among loci classified as outliers in the *F*_ST_ outlier scan (na – non-applicable).

The coupling between ancestry and environmental associations was particularly clear for the turbot, where the environmental outliers were consistently associated with the highly differentiated loci and found across most of the genome (Figure 3). However, this coupling was less evident in other species, where some loci were more strongly associated with the environment than the average genetic marker, but weakly associated with ancestry and demographic history. For example, both plaice and dab showed environmental outliers localized on specific regions along the chromosomes (chromosomes 7, 12, 22, 23 for dab, and 14, 19, 21, 23 for plaice) while other differentiated genomic regions were not associated with the environmental cline (Figure 3). Such data suggest a decoupling of signals associated with ancestry and environment, possibly linked to environmental adaptation. Moreover, for the pelagic flounder, only a single highly differentiated and environment-associated locus was detected, which could represent a strong candidate for local adaptation. Nevertheless, this decoupling could also reflect the complex interplay of gene flow and selection at sites linked to incompatibilities in the later phases of a secondary contact (Gagnaire *et al.*, 2015).

We also found evidence for the effects of selection based on the spatial distribution of the derived mutations at outlier loci (Table 2 and JAFS from Figure 3). Although this was again quite variable across species, we did find an increase in the frequency of the derived allele in the Baltic Sea for more than 60% of the *F*_ST_ outlier loci (80% of the top 0.1% outliers) for the Baltic Sea turbot and the DEM Baltic Sea flounder (Table 2 and JAFS from Figure 3d,e). This disequilibrium is quite unexpected under a pure demographic scenario of secondary contact (which should affect both ancestral and derived alleles randomly through drift) and therefore suggest that Baltic Sea turbot and DEM flounder samples carry stronger signals from non-random evolutionary pressures. Two main processes of selection could result in this disequilibrium, both acting preferentially on the Baltic Sea lineages.

Firstly, background selection can explain this pattern, in particular in regions of the genome that experience low recombination (Perrier and Charmantier, 2018). This pattern was observed for the highest peak of *F*_ST_ in turbot (Figure 3), which is located in a genomic region of low recombination coinciding with the location of the centromere of chromosome 1 (Maroso *et al.*, 2018; Martínez *et al.*, 2008). This region also showed reduced diversity in both populations leading to low *d*_XY_ (Figure S11), which is typically expected from the long-term effect of background selection (Cruickshank and Hahn, 2014). Moreover, the BS effective population size was inferred to be three times smaller than that of the NS in this species. The reduction in effective population size of the BS lineages could have favoured the accumulation of more deleterious mutations during the isolation phase, and therefore increased the signature of background selection in the region of low recombination rates (Gagnaire *et al.*, 2018; Roesti *et al.*, 2013). Then, the current resistance to gene flow could have protected the differential signature of background selection over the period of secondary contact (Duranton *et al.*, 2018).

Secondly, the higher proportions of derived outlier mutations in the BS could also be linked to directional selection associated with adaptation to the brackish environment. As this effect was detected only in the two species with the longest isolation phase, it is possible that the current Baltic Sea turbot and DEM Baltic Sea flounder were isolated in a brackish environment during the LGM. In this case, their isolation may have facilitated the spread of new adaptive mutations through the suppression of migration load (Lenormand, 2002; Yeaman, 2015). Although the Baltic basin was entirely covered by ice during the LGM, marine water was captured within the present day North Sea (near sampling sites 2-3 in Figure 1) during the LGM (Willmes *et al.*, 2016, Kettle *et al.*, 2011), which could potentially correspond to a brackish refugia located near the present day Baltic Sea. Further studies using ancient DNA from sediments could provide data to test this hypothesis. Background selection and local adaptation are not mutually exclusive processes and could thus have contributed together to the observed excess of the derived mutations in the Baltic Sea.

In contrast to the patterns observed for the highly differentiated species, we did not find a clear pattern in the distribution of the derived mutations in the species more permeable to gene flow (pelagic flounder, dab and plaice). In these species, there was a slight tendency towards fewer derived mutations in the Baltic Sea for the top 5% *F*_ST_ outliers. This deficit in derived mutations could reflect the loss of diversity associated with a recent colonization of the region across the environmental gradient, and a small increase of the derived alleles among the top 0.1% *F*_ST_ outliers, which could reflect recent *de novo* adaptation (Johannesson and André, 2006). Altogether, these findings thus suggest that the relative contribution of colonization events, local adaptation and ancient demographic history appears to be species-dependent, regardless the fact that all species occur in the same environmental gradient.

## Conclusion

This study illustrates the great diversity of genetic signatures associated with the divergence of populations of marine fishes in the Baltic Sea. It is generally assumed that the diversification process involved during the colonization of new environment starts from ancestral populations at equilibrium (Momigliano *et al.*, 2017). The replicates of population divergence analysed here suggest that this equilibrium was not met during the colonization of the Baltic Sea, and that the divergence of populations predated (i.e. five to ten times older) the access to a Baltic Sea habitat, approximately 8 000 years ago (Björck, 1995). Consequently, the flatfish populations currently inhabiting the Baltic Sea and the North Sea may have been isolated in different glacial refugia during the LGM. Two different periods of secondary contact were estimated, which may then suggest the existence of more than two refugia. The plaice was the only species in which a scenario of sympatric divergence was supported. Here, divergence may have been facilitated by the presence of two large structural variants resistant to gene flow (more details in Le Moan *et al.*, 2019). Importantly, the genomic regions inferred to be under divergent selection were not shared across species. These results contrast with reported evidence of genomic convergence observed in parallel evolution (Jones *et al.*, 2012; Ravinet *et al.*, 2016), and evolutionary radiations (Hench *et al.*, 2019; Muschick *et al.*, 2012; Stryjewski and Sorenson, 2017). Our results suggest that recent and independent replicates of ecological divergence resulted in limited convergence due to the absence of shared genetic variation, complex and species-specific genomic architecture of traits under selection, and the recent timing of divergence, which may have prevented background selection to lead to similar signatures of selection (Burri, 2017). Thus, adaptation to the environmental gradient in these species is likely to have relied on independent genomic pathways, highlighting the importance of genomic background as raw material for the action of selection in natural and genetically diverse populations (Blount *et al.*, 2018).

## Methods

### Sampling

In order to study replicates of ecological speciation events, we sampled four flatfish species that have successfully colonized the environmental gradient of the Baltic Sea. The species belong to two distant families, the Scophthalmidae: turbot *(Scophthalmus maximus),* and the Pleuronectidae: common dab *(Limanda limanda),* European plaice *(Pleuronectes plastessa),* and European flounder *(Platichthys flesus).* The flounder was recently described as two different species, *P. flesus* and *P. solemdali* (Momigliano *et al.*, 2018), corresponding to two different spawning strategies, respectively pelagic and demersal. We therefore considered these two species as ecotypes of the same species (PEL and DEM) for the purpose of this study. For each species, 25 individuals were sampled from six to 12 sampling sites in the Atlantic Ocean and the Baltic Sea (Figure 1a). The majority of samples were collected during the spawning season of 2016 and 2017. For the northern sampling site of the Baltic Sea (sampling site 12), we included a few turbot samples from Nielsen et al. (2004) and flounder samples from Hemmer-Hansen *et al.* (2007). Moreover, we included four additional turbot sampling sites within the North Sea and the English Channel (sampling sites 1 to 4) collected by Vandamme *et al.* (2014). We additionally sampled eight North Sea brill *(Scophthalmus rhombus)* individuals, a closely related species to the turbot, to polarise the turbot polymorphism for the demographic inferences and the selection tests. In total, 860 Individuals were sampled for the purpose of this study, and further details about the sampling strategy can be found in Table S9.

### Library preparation and sequencing

Genomic DNA was extracted from either gills or fins using the DNeasy Blood Tissue kit (Qiagen). DNA concentration was measured using the Broad Range protocol of the Qubit version 2.0^®^ and standardized to 20 ng/μl. Fifteen double-digestion (dd-RAD) libraries were constructed by randomly pooling between 60 and 75 barcoded samples from various locations and species, following a modified version of Poland and Rife (2012). This protocol involved the Pst1 and Msp1 restriction enzymes with low and high cut site frequencies, respectively. Size selection was performed in two steps in order to conserve sequences between 350 and 500 bp. The first step was done after the pooling of the barcoded samples on agarose gels, and the second with AMPure^®^ beads after the PCR amplification (14 cycles). The quality of the size selection was assessed on a Bioanalyzer 2100 using the High Sensitivity protocol (Agilent Technologies). Each library with a consistent size selection was sequenced for paired-end on one Illumina HiSeq4000 lane (2*101 bp) by BGI TECH SOLUTIONS (HONGKONG) CO.

### Bioinformatics

The libraries were processed using the “ref-map” pipeline from Stacks version 1.46 (Catchen *et al*., 2013). The pooled sequences were demultiplexed by barcode using the “process radtag” program set to keep only sequence with phred33 quality above 10, and were trimmed to 85 bp using trimmomatic (Bolger *et al.*, 2014). On average, we obtained six million reads per sample (Figure S12). The reads were aligned to a reference genome using bwa (Li and Durbin, 2009). Both turbot and brill samples were aligned to the turbot (*S. maximus)* genome (Figueras *et al.*, 2016; Maroso *et al.*, 2018), with an average of 99% and 98% of reads properly mapped (Figure S12). The sequences of the three remaining species were mapped against the genome of the Japanese flounder, *Paralichthys olivaceus* (Shao *et al.*, 2017) with an average 70% of the reads mapped for dab, 68% for flounder and 60% for the plaice (Figure S12). The aligned sequences were processed using the “pstacks” programme with a minimum coverage (m) of 5X within one individual to consider a stack of reads as consistent biological sequences. Stacks mapping to the same position in the genome were considered as alleles of one locus. These alleles were stored in two independent catalogues, one for each reference genome, with the “cstacks” programme. Subsequently, every sample was mapped back to the catalogues and genotyped for all the polymorphic sites using “pstacks”. The average coverage before filtration was 12X (Figure S12). Only bi-allelic SNPs present in at least 80% of the individuals within each sampling site and with a maximum heterozygosity of 0.80 were called using the population programme. We removed individuals with more than 10% missing data. Finally, we removed singletons and SNPs with a significant departure from Hardy-Weinberg Equilibrium proportions (p-value 0.05) in more than 60% of the sampling sites using vcftools (Danecek *et al.*, 2011). The average coverage after these filtration steps was 55X (Figure S12). Two datasets were then constructed per species, one to analyse the population structure along the NBTZ and the second one to infer the demographic history of the Baltic Sea colonization. We used all the sampling sites for the “population structure” datasets and only the two most distant sampling sites and 25 random samples of an outgroup species (detailed table S7) for the “demographic inference” dataset. Thus, we used a total of eight datasets (two per species) for the purpose of this study.

### Population structure

The genetic structure was assessed independently for each species using the “population structure” dataset. We kept SNPs with a minor allele frequency above 0.05 and only one random SNP per bin of 1 kb along the chromosome to limit effects from physical LD (Figure S13) and to fulfil the requirements of several software used in the study. In total, we used 3 348 SNPs for turbot (0.91 % of total missing data), 5 472 SNPs for flounder (0.68%), 6 685 for plaice and 3 468 for dab (1.54%). The genetic variation within each dataset was visualized using principal component analyses (PCAs) from the R package adegenet (Jombart, 2008). Then, the most likely number of populations and ancestry of each individual were analyzed using the R package ConStruct (Bradburd *et al.*, 2018), by setting the appropriate number of clusters (k) based on AIC tests. We used k = 2 in all cases, except for the flounder which was performed on k = 2, k = 3, and k = 2 but without including the DEM individuals (Figure S2). In order to map the population shifts between the North Sea and the Baltic Sea, we fitted a cline ancestry analysis for each run of k = 2. More precisely, we used a generalized linear model with a binomial family to fit the probability of each fish belonging to the Baltic Sea cluster as a function of the distances of these fish from the North Sea (distance, species, and their interaction was set as fixed parameters in the model). The effect of each parameter was then tested in a top-down approach based on likelihood ratio tests. The presence of potential hybrids between the different genetic clusters was assessed by running the “snapclust” function in adegenet (Beugin *et al.*, 2018). We estimated the Weir and Cockerham (1984) pairwise *F*_ST_ between sampling sites and tested for significant genetic differences with 1 000 permutations over loci using the R package StAMPP (Pembleton *et al.*, 2013). Finally, we used the approach applied in Souissi et al. (2018) to evaluate geographical patterns of allelic clines along the transition zone at every locus without pruning the datasets for physical LD. Specifically, we used the R package HZAR (Derryberry *et al.*, 2014) to calculate the center and the slope of the allelic cline for each SNP. We then represented the estimate of the slopes depending of their centers across the NBTZ using the R package ggplot2 (Wickham and Winston, 2008).

### Environmental data

The environmental data for salinity, sea surface temperature and temperature at the bottom were downloaded from the Global Ocean 1/12° Physics Analysis and Forecast (http://marine.copernicus.eu/), updated daily over a period of 3 months during the spawning season of the study species (February to April 2016). The environmental data for each site were obtained by averaging the measurement over 50 km^2^ around the sampling location (Table S9) during the entire period extracted. Additionally, we applied a smoothing function using the R package Stats to obtain the prediction of salinity for each allele frequency cline center estimate.

### Genomic architecture of differentiation

We kept the two most distant sampling sites of the “population structure” dataset to analyse samples located outside the NBTZ and limit the consequences of recent hybridization between genetic clusters on the genomic architecture of population differentiation. These datasets were not filtered for LD or for minor allele frequency. For each SNP, we computed the genetic diversity π per population using the software vcftools (Danecek *et al.*, 2011), and the pairwise *F*_ST_ between populations (Hudson *et al.*, 1992) using the R package KRIS (Chaichoompu *et al.*, 2018). Using a custom R script (which should be provided), we calculated the between populations diversity *d*_XY_ as explained in Cruickshank and Hahn (2014). The three statistics, π, *d*_XY_ and *F*_ST_ were averaged over sliding windows of 100 kbp. These averaged statistics were then standardized, i.e. multiplied by the number of polymorphic SNPs and divided by the sequence length in the window. Finally, we estimated Spearman rank correlation coefficients between species for the standardized statistics to assess similarity in the patterns of genetic diversity and NS-BS divergence. Here, we only used the three species of Pleuronectidae (plaice, flounder and dab) that were aligned to the same reference genome.

### Demographic inferences

The demographic inferences were performed using the “demographic inference” dataset (see bioinformatics part). We retained only one SNP per 1 kb to limit effects from physical LD (Figure S13). To control for high LD within the putative structural variants in the plaice genome, we ran an additional comparison including only one highly differentiated SNP within the linkage block. The results did not change significantly (i.e. the same models were supported) and we report only the inferences without high LD. Demographic inferences were performed using a composite likelihood approach implemented in the modified version of δaδi (Gutenkunst *et al.*, 2010) from Tine et al. (2014). This software infers the demographic history from contrasted models of evolution using the Joint Allele Frequency Spectrum (JAFS). In our analyses, the data fit to 11 models (details of the models in Rougeux *et al.* 2017) were compared for each species. Specifically, we compared models of strict isolation (SI) to models including migration. This migration was either continuous along the process of divergence (IM=isolation with migration) or discontinuous, either in the beginning of the divergence (AM=ancestral migration) or after an isolation phase during secondary contact (SC). In order to test the effect of heterogeneity of recombination rates leading to variable effective size along the chromosome, we included the possibility of reduced effective population size (_r_N_E_) to a certain proportion of loci to the models (SI2N, IM2N, AM2N and SC2N). Similarly, to test for effects of selection leading to heterogeneous barriers to gene flow between populations undergoing divergence, we included a reduced migration rate (rm) to a certain proportion of loci in the models (IM2m, AM2m, and SC2m). To improve the predictions of the model, we used the unfolded JAFS with genotypes of few individuals from an outgroup species (outgroup detailed Table S7) to identify the derived allele at each SNP (allele not fixed in the outgroup). To ensure correct unfolding of the spectrum, we only used SNPs that were polymorphic within the focal pair of populations and fixed in the outgroup species for these analyses. Therefore, SNPs that were differentially fixed in the species or polymorphic in both species were removed. Singletons were not considered for the inferences and were hidden on the JAFS using the -z parameter in δaδi. Each model was run independently 30 times for a total of 1 980 runs. The predictions of each model were then compared based on the goodness of fit using AIC. We also calculated the weighted AIC (wAIC) as presented in Rougeux *et al.* (2017) reflecting the predictability of a given model relative to the other models. A high value of wAIC (0.5 – 1) corresponds to the best model. An AIC difference above 10 between the best and the second best model leads to a wAIC of 1 which represents a strong support for the corresponding demographic scenario. Only the best models for each species are shown in Table 1, the other models are reported in the supplementary material (Table S7). To finish, the parameters estimated form the model relatively to θ were transformed into biological numbers following the explanations provided in the user manual of δaδi (Gutenkunst *et al.*, 2010). These transformations were performed using a generation time of 3.5 years (Erlandsson *et al.*, 2017) and a mutation rate of 10^-8^ per generation (The 1000 Genomes Project Consortium, 2010).

### Test for selection

To identify SNPs as candidates for carrying signals of selection, we applied two different methods, which are able to take the complex demographic history of the Baltic Sea populations into account. Firstly, we used the software Bayenv over the six sampling sites repeated across the four species (from sites 5 to 10, Figure 1) to detect SNPs with allelic frequencies more associated to the environmental gradient than to genome wide structure. The population covariance matrix and environmental associations for individual loci were estimated through 100000 iterations. Three independent runs with unique random number seeds were performed and only SNPs with a Bayes Factor above one in all three runs were considered as good outlier candidates. Secondly, we used the results from the demographic inferences, applying only the neutral parameters of divergence, to simulate the differentiation for 100 000 independent neutral loci between the two most distant sampling sites, using different combinations of missing data, with the coalescent simulator msms (Ewing and Hermisson, 2010). From the simulated dataset, the *F*_ST_ between populations and the overall expected heterozygosity were calculated for all simulated loci using the software msstat (Thornton, 2003). Then, we estimated the neutral envelope of differentiation following the approach of Beaumont and Nichols (1996) by fitting a quantile regression to the 95, 99, and 99.9% quantile over bins of 0.02 of expected heterozygosity. Using the quantile regression models, we predicted the maximum value of *F*_ST_ for 5%, 1% and 0.1% upper quantiles depending of the observed heterozygosity of each SNP in the datasets. All the loci with an observed value of *F*_ST_ above the predicted value of *F*_ST_ quantiles were considered as outliers. Finally, from the loci identified as *F*_ST_ outliers, we estimated the proportion of loci with the derived allele increasing in frequency in the Baltic Sea. To do this, we used only loci sequenced in both focal and outgroup species and polarized the alleles following the procedure described for the generation of the JAFS (see above).

## Acknowledgement

We gratefully thanks Dorte Meldrup for her assistance in the library preparation and well as Sara Vandamme and Filip Volckaert for the providing of the turbot samples from the North Sea. We also thanks Henrik Baktoft for his help regarding the statistical model and Andres Blanco his help with the sharing of the Turbot genome. This study received financial support from The European Regional Development Fund (Interreg V-A, project “MarGen”), from Ørsted Foundation and from the Otto Mønsteds fond.

## Supplementary material

### Population structure

**Figure S1:**
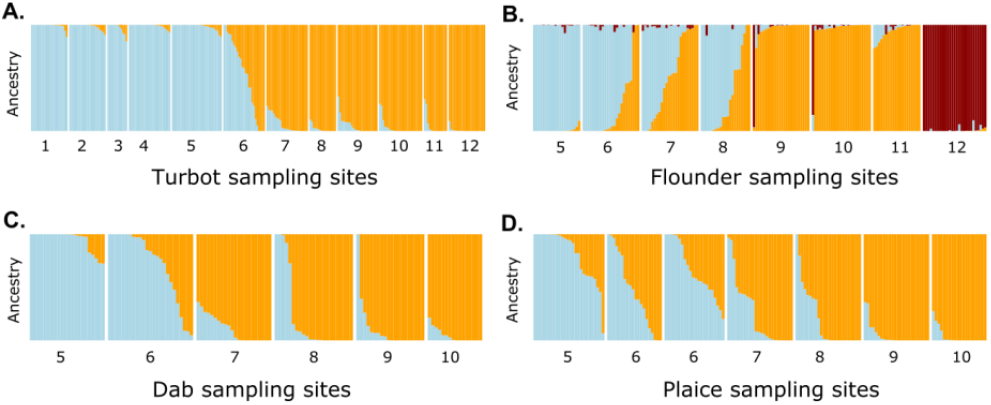
Structure plot from Construct analyses for the appropriate number of clusters based on AIC selection for the samples of A) turbot, B) flounder (PEL and DEM), C) dab and D) plaice.

**Figure S2:**
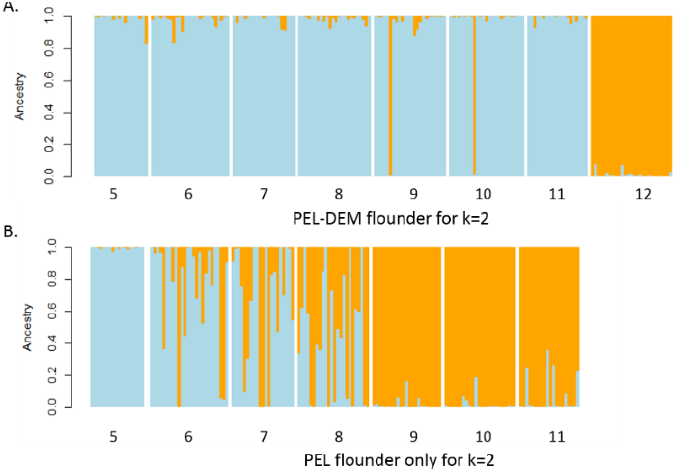
Structure plot from Construct analyses for k=2 in the flounder samples, with A) both pelagic and demersal ecotypes, and B) the pelagic ecotypes alone.

**Table S1:** Estimates of the parameter from the generalised linear model with binomial family for the variation of estimated Baltic Sea ancestry as a function of geographical distance, the species, and the interaction between the two factors (all significant: Chi^2^ = 19.44, p-value = 0.0006, df = 4)

| Factors | Estimate | z-value | Std Error | Pr(>\|z\|) |
| --- | --- | --- | --- | --- |
| (Intercept) | -6.4592 | 1.8808 | -3.4340 | 0.0006*** |
| Distance | 0.0096 | 0.0025 | 3.9000 | 0.0001*** |
| Flounder PEL-PEL | -3.6599 | 2.9566 | -1.2380 | 0.2158 |
| Flounder PEL-DEM | -6.5043 | 2.7641 | -2.3530 | 0.0186* |
| Plaice | 4.0242 | 1.9448 | 2.0690 | 0.0385* |
| Turbot | 1.1805 | 2.1752 | 0.5430 | 0.5873 |
| Distance : Flounder PEL | 0.0037 | 0.0038 | 0.9680 | 0.3328 |
| Distance : Flounder PEL-DEM | 0.0007 | 0.0030 | 0.2260 | 0.8214 |
| Distance : Plaice | -0.0046 | 0.0026 | -1.7790 | 0.0753 |
| Distance : Turbot | -0.0008 | 0.0029 | -0.2820 | 0.7783 |

**Figure S3:**
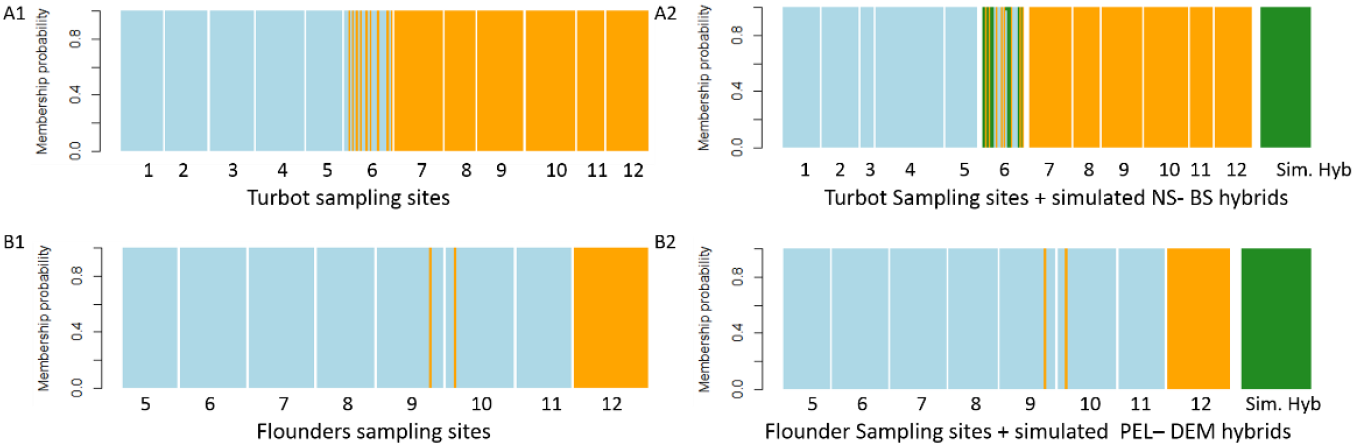
Structure plot from the snapclust function of adegenet without hybrid detection (A1 and B1) and with hybrids detection (A2 and B2) for the turbot (A) and the two ecotypes (PEL and DEM) of flounder (B). Yellow bars correspond to samples assigned to the Baltic Sea turbot and to the demersal ecotype of flounder. Blue corresponds to North Sea turbot and pelagic flounder. Green corresponds to individuals classified as hybrids, where “Sim Hyb.” (A2 and B2) in both species correspond to 30 simulated hybrids between the two most distant sampling sites.

**Figure S4:**
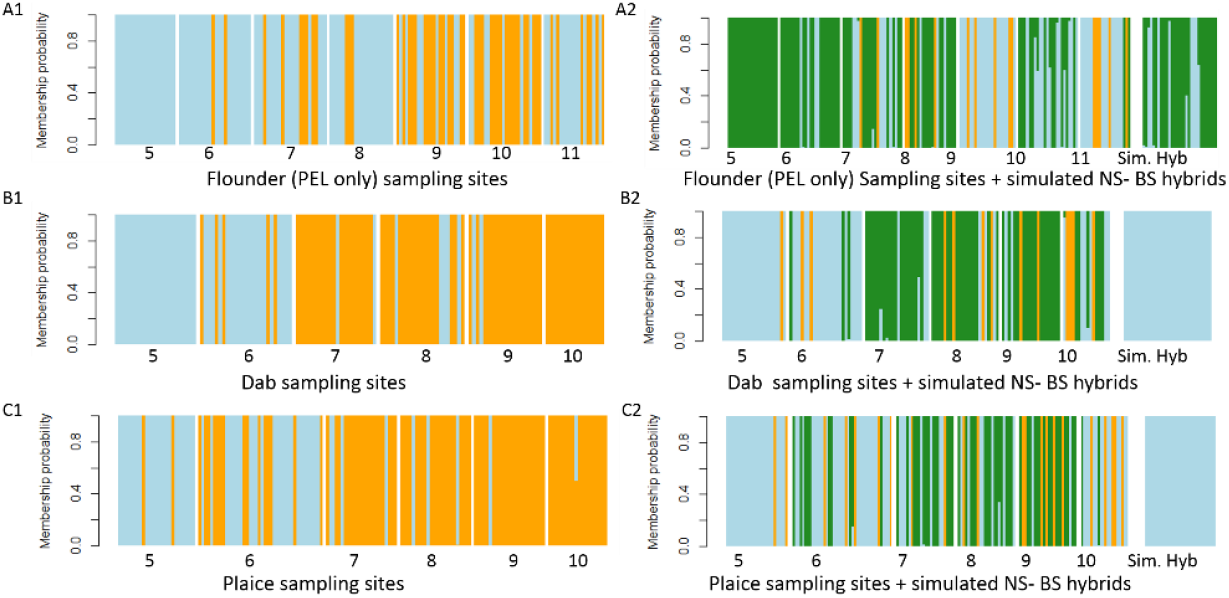
Structure plot from the snapclust function of adegenet without hybrid detection (A1, B1 and C1) and with hybrid detection (A2, B2 and C2) for the pelagic flounder only (A), the dab (B) and the plaice (C). Yellow bars correspond to samples assigned to the Baltic Sea lineages, blue to North Sea lineages, and green to putative hybrids (A2, B2 and C2). The “Sim Hyb.” (right) correspond to 30 simulated hybrids between the two most distant sampling sites.

**Figure S5:**
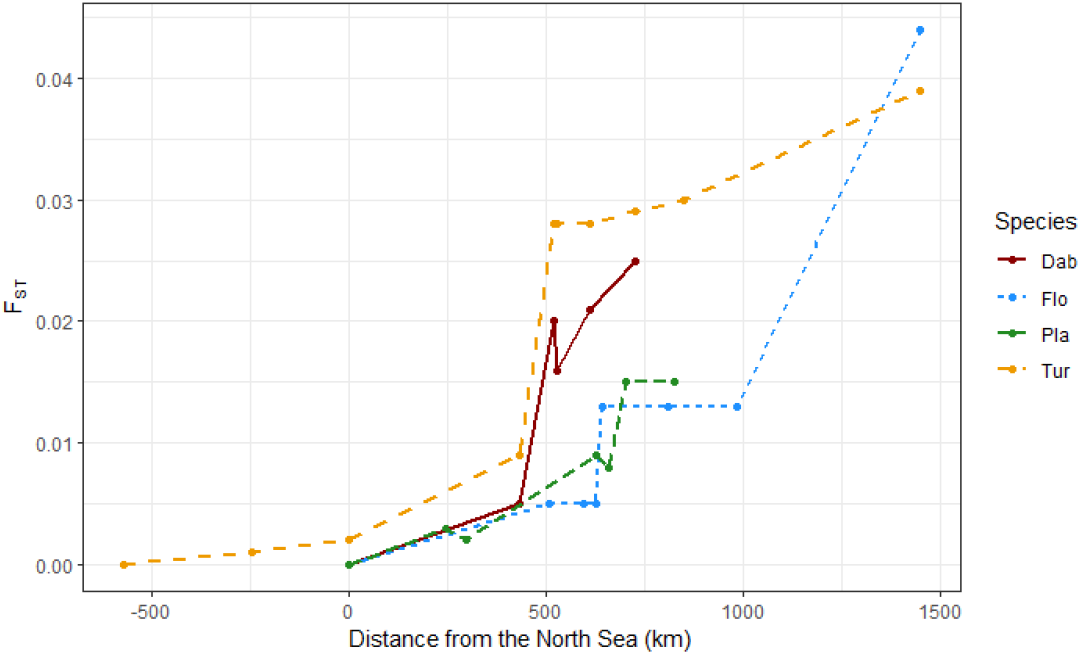
Pairwise *F*_ST_ by species, for sampling site 5 in the North Sea compared with other sampling sites as a function of the distance to the North Sea (site 5 in Figure 1). The two dots with negative value are the site from the English Channel from which only turbot were collected

**Table S2:**
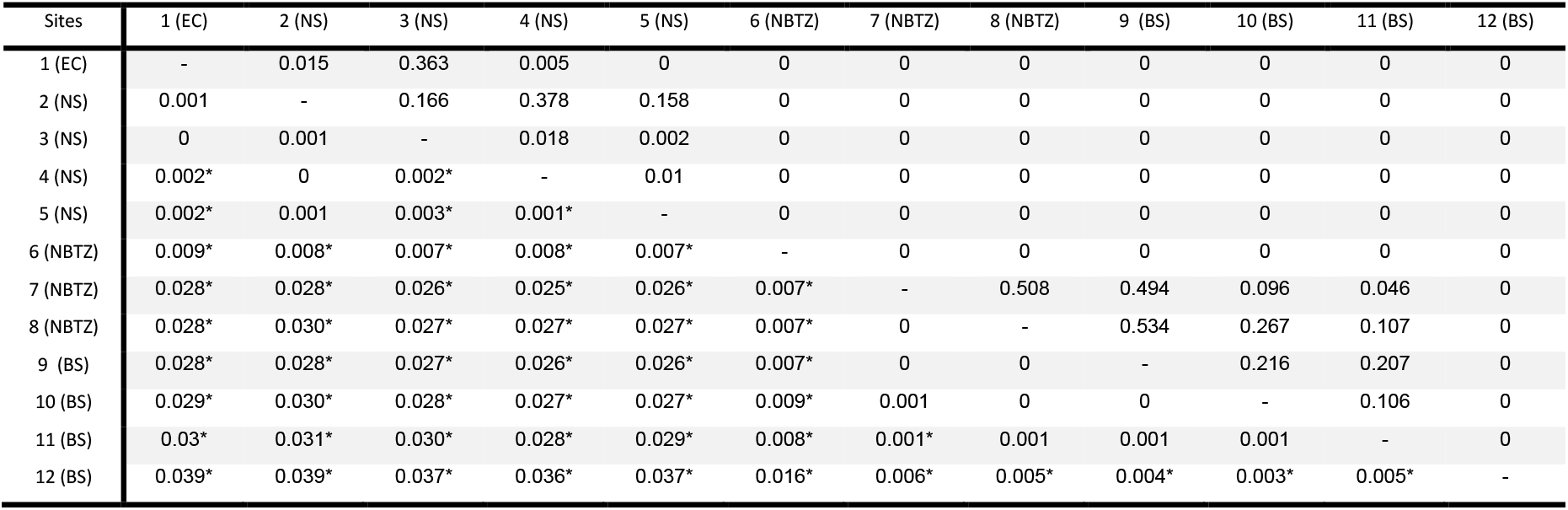
Pairwise Fst between turbot populations, where values bellow the diagonal are Fst estimates and values above the diagonal show the p-value of the permutation test.

| Sites | 1 (EC) | 2 (NS) | 3 (NS) | 4 (NS) | 5 (NS) | 6 (NBTZ) | 7 (NBTZ) | 8 (NBTZ) | 9 (BS) | 10 (BS) | 11 (BS) | 12 (BS) |
| --- | --- | --- | --- | --- | --- | --- | --- | --- | --- | --- | --- | --- |
| 1 (EC) | - | 0.015 | 0.363 | 0.005 | 0 | 0 | 0 | 0 | 0 | 0 | 0 | 0 |
| 2 (NS) | 0.001 | - | 0.166 | 0.378 | 0.158 | 0 | 0 | 0 | 0 | 0 | 0 | 0 |
| 3 (NS) | 0 | 0.001 | - | 0.018 | 0.002 | 0 | 0 | 0 | 0 | 0 | 0 | 0 |
| 4 (NS) | 0.002* | 0 | 0.002* | - | 0.01 | 0 | 0 | 0 | 0 | 0 | 0 | 0 |
| 5 (NS) | 0.002* | 0.001 | 0.003* | 0.001* | - | 0 | 0 | 0 | 0 | 0 | 0 | 0 |
| 6 (NBTZ) | 0.009* | 0.008* | 0.007* | 0.008* | 0.007* | - | 0 | 0 | 0 | 0 | 0 | 0 |
| 7 (NBTZ) | 0.028* | 0.028* | 0.026* | 0.025* | 0.026* | 0.007* | - | 0.508 | 0.494 | 0.096 | 0.046 | 0 |
| 8 (NBTZ) | 0.028* | 0.030* | 0.027* | 0.027* | 0.027* | 0.007* | 0 | - | 0.534 | 0.267 | 0.107 | 0 |
| 9 (BS) | 0.028* | 0.028* | 0.027* | 0.026* | 0.026* | 0.007* | 0 | 0 | - | 0.216 | 0.207 | 0 |
| 10 (BS) | 0.029* | 0.030* | 0.028* | 0.027* | 0.027* | 0.009* | 0.001 | 0 | 0 | - | 0.106 | 0 |
| 11 (BS) | 0.03* | 0.031* | 0.030* | 0.028* | 0.029* | 0.008* | 0.001* | 0.001 | 0.001 | 0.001 | - | 0 |
| 12 (BS) | 0.039* | 0.039* | 0.037* | 0.036* | 0.037* | 0.016* | 0.006* | 0.005* | 0.004* | 0.003* | 0.005* | - |

**Table S3:** Pairwise Fst for flounder populations (below diagonal) and p-value of permutation test (above diagonal).

| Sites | 5 (NS) | 6 (NBTZ) | 7 (NBTZ) | 8 (NBTZ) | 9 (BS) | 10 (BS) | 11 (BS) | 12 (DEM) |
| --- | --- | --- | --- | --- | --- | --- | --- | --- |
| 5 (NS) | - | 0 | 0 | 0 | 0 | 0 | 0 | 0 |
| 6 (NBTZ) | 0.005* | - | 0 | 0.291 | 0 | 0 | 0 | 0 |
| 7 (NBTZ) | 0.005* | 0.002 | - | 0 | 0 | 0 | 0 | 0 |
| 8 (NBTZ) | 0.005* | 0 | 0.003* | - | 0 | 0 | 0 | 0 |
| 9 (BS) | 0.013* | 0.007* | 0.003* | 0.005* | - | 0.003 | 0 | 0 |
| 10 (BS) | 0.013* | 0.007* | 0.002* | 0.005* | 0.001* | - | 0 | 0 |
| 11 (BS) | 0.013* | 0.007* | 0.005* | 0.005* | 0.004* | 0.004* | - | 0 |
| 12 (DEM) | 0.044* | 0.045* | 0.042* | 0.044* | 0.043* | 0.043* | 0.048* | - |

**Table S4:**
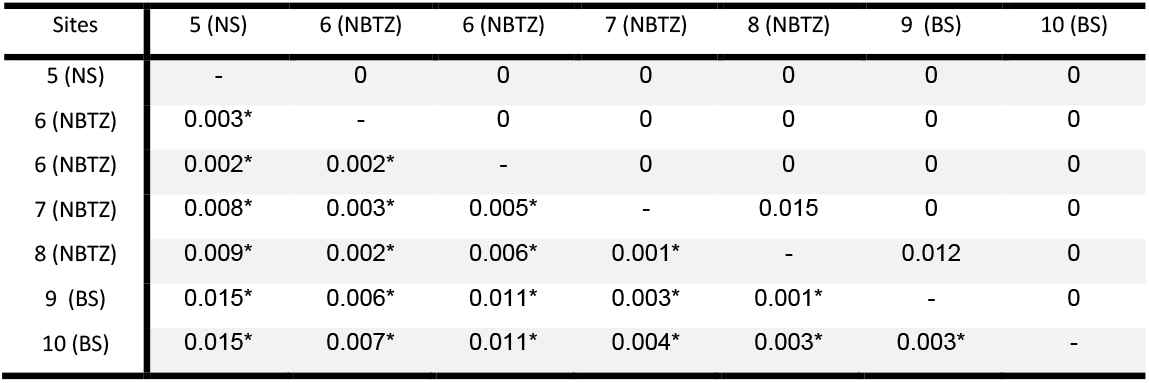
Pairwise Fst for plaice populations (below diagonal) and p-value of permutation test (above diagonal).

| Sites | 5 (NS) | 6 (NBTZ) | 6 (NBTZ) | 7 (NBTZ) | 8 (NBTZ) | 9 (BS) | 10 (BS) |
| --- | --- | --- | --- | --- | --- | --- | --- |
| 5 (NS) | - | 0 | 0 | 0 | 0 | 0 | 0 |
| 6 (NBTZ) | 0.003* | - | 0 | 0 | 0 | 0 | 0 |
| 6 (NBTZ) | 0.002* | 0.002* | - | 0 | 0 | 0 | 0 |
| 7 (NBTZ) | 0.008* | 0.003* | 0.005* | - | 0.015 | 0 | 0 |
| 8 (NBTZ) | 0.009* | 0.002* | 0.006* | 0.001* | - | 0.012 | 0 |
| 9 (BS) | 0.015* | 0.006* | 0.011* | 0.003* | 0.001* | - | 0 |
| 10 (BS) | 0.015* | 0.007* | 0.011* | 0.004* | 0.003* | 0.003* | - |

**Table S5:** Pairwise Fst for dab populations (below diagonal) and p-value of permutation test (above diagonal).

| Sites | 5 (NS) | 6 (NBTZ) | 7 (NBTZ) | 8 (NBTZ) | 9 (BS) | 10 (BS) |
| --- | --- | --- | --- | --- | --- | --- |
| 5 (NS) | - | 0 | 0 | 0 | 0 | 0 |
| 6 (NBTZ) | 0.005* | - | 0 | 0 | 0 | 0 |
| 7 (NBTZ) | 0.020* | 0.012* | - | 0 | 0.292 | 0.015 |
| 8 (NBTZ) | 0.016* | 0.010* | 0.002* | - | 0 | 0 |
| 9 (BS) | 0.021* | 0.013* | 0.000* | 0.004* | - | 0.001 |
| 10 (BS) | 0.025* | 0.017* | 0.002* | 0.005* | 0.004* | - |

**Figure S6:**
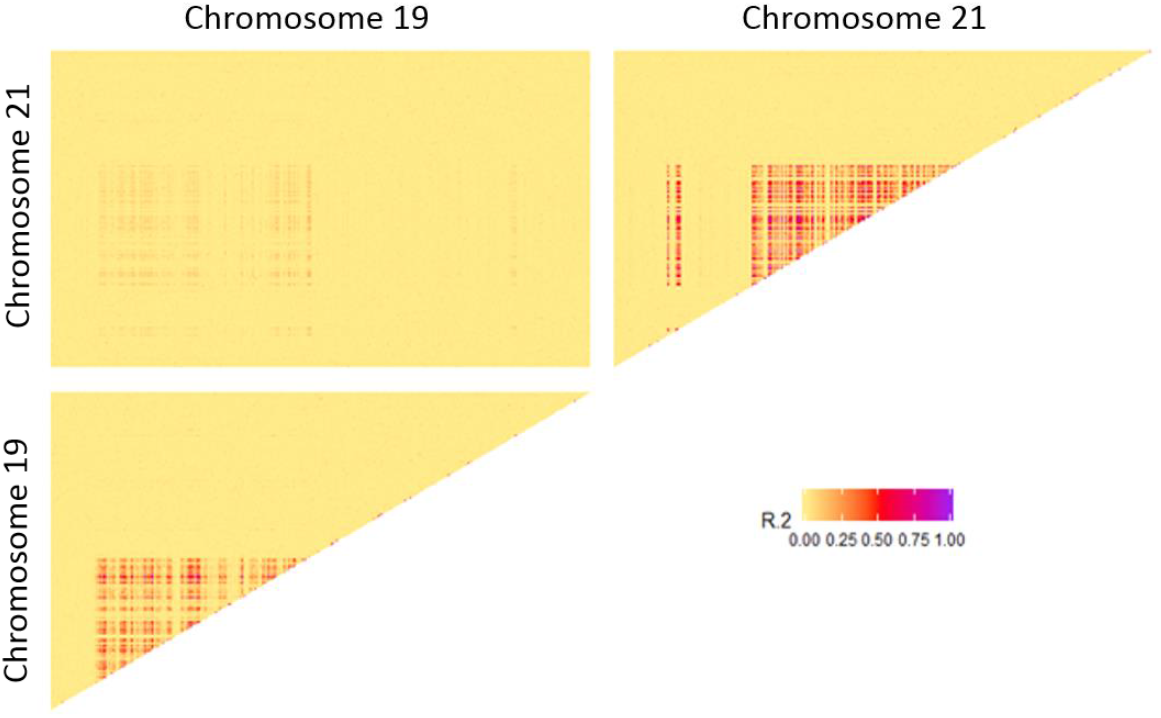
LD-heatmap from the plaice dataset showing the linkage disequilibrium between all pairs of loci from chromosomes 19 and 21 carrying putative structural variants. The regions with LD above 0.5 cover more than 8Mbp along each chromosome.

**Table S6:**
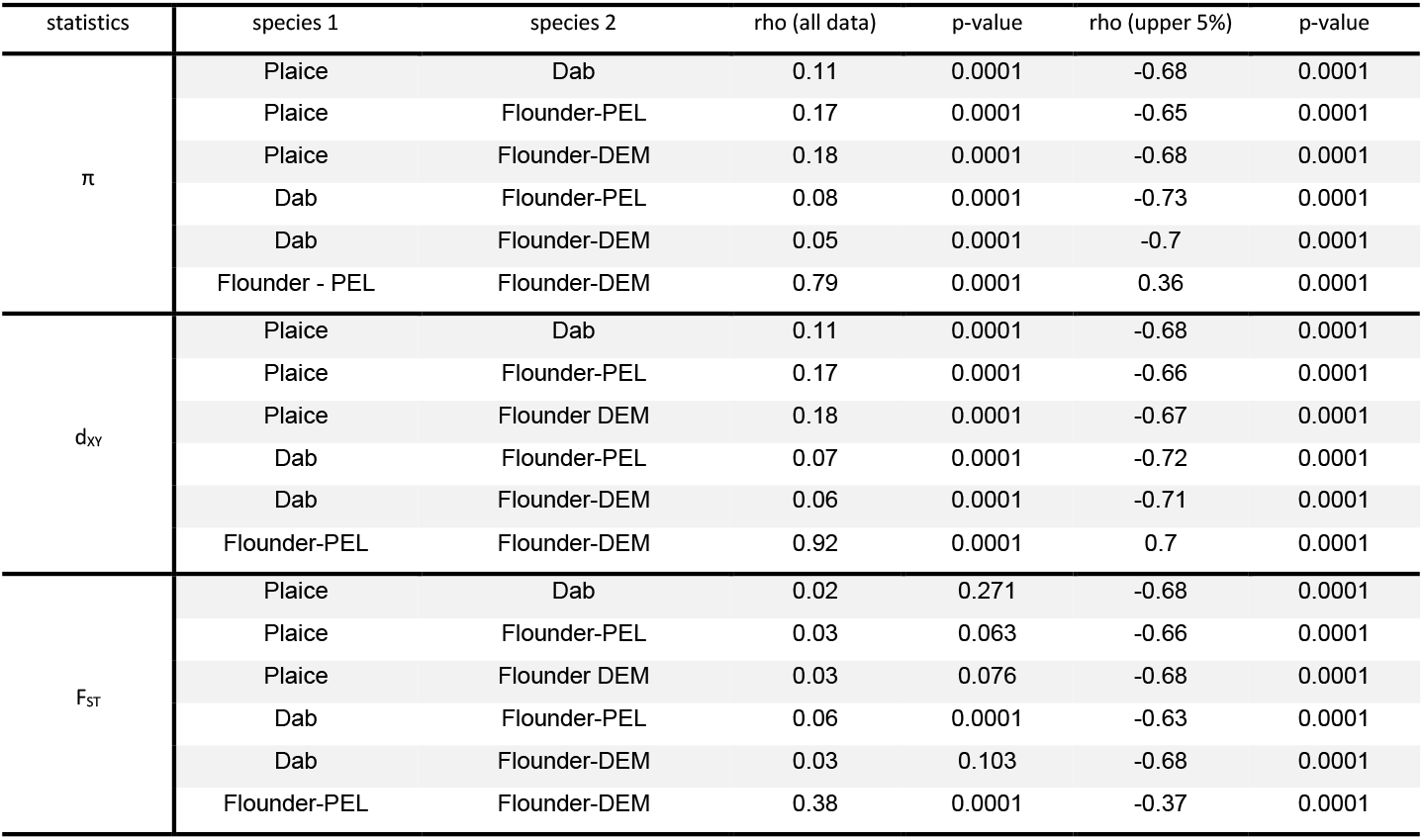
Spearman correlation (rho) between the statistic of diversity and divergence calculated for different pairs of species aligned to the same genome in a sliding windows approach (bin of 100kb) on the entire dataset (all) on the upper quantile (5%) of each statistics in each species

### Demographic inference

**Table S7:**
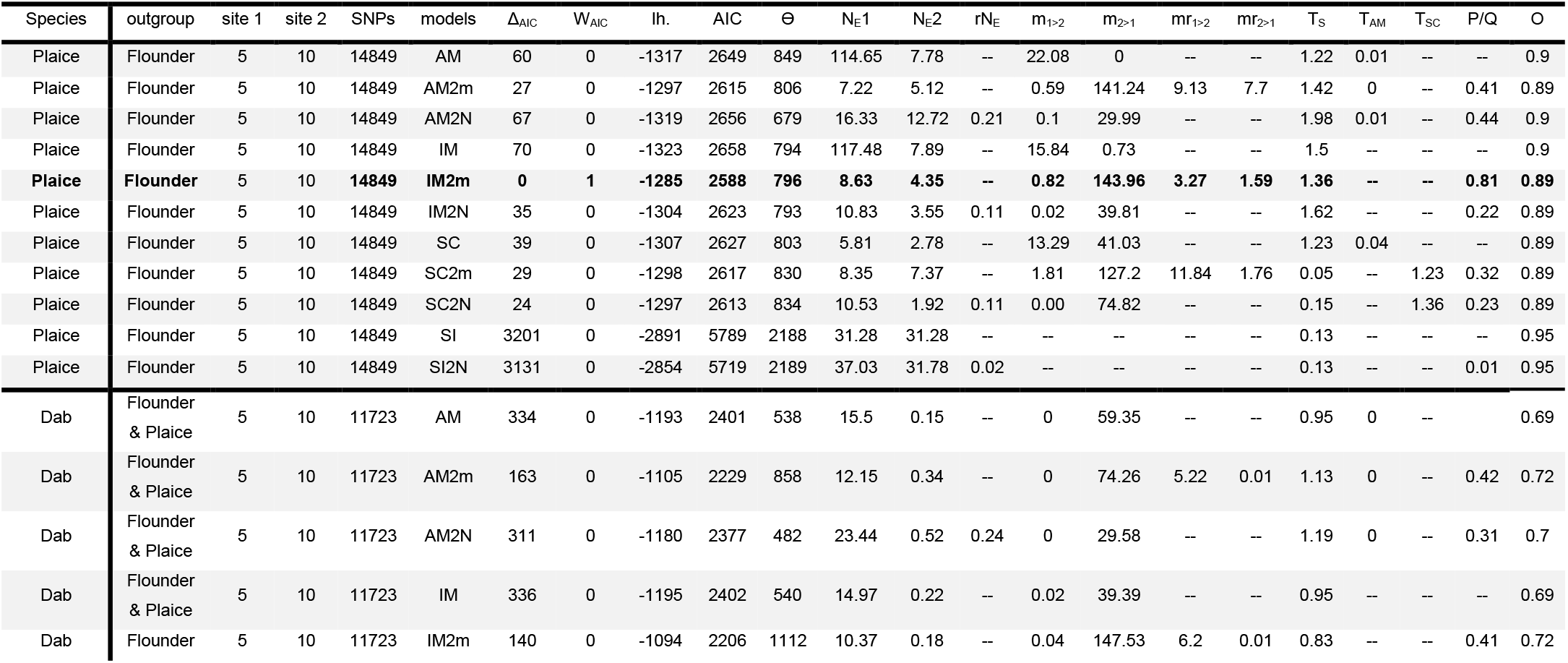
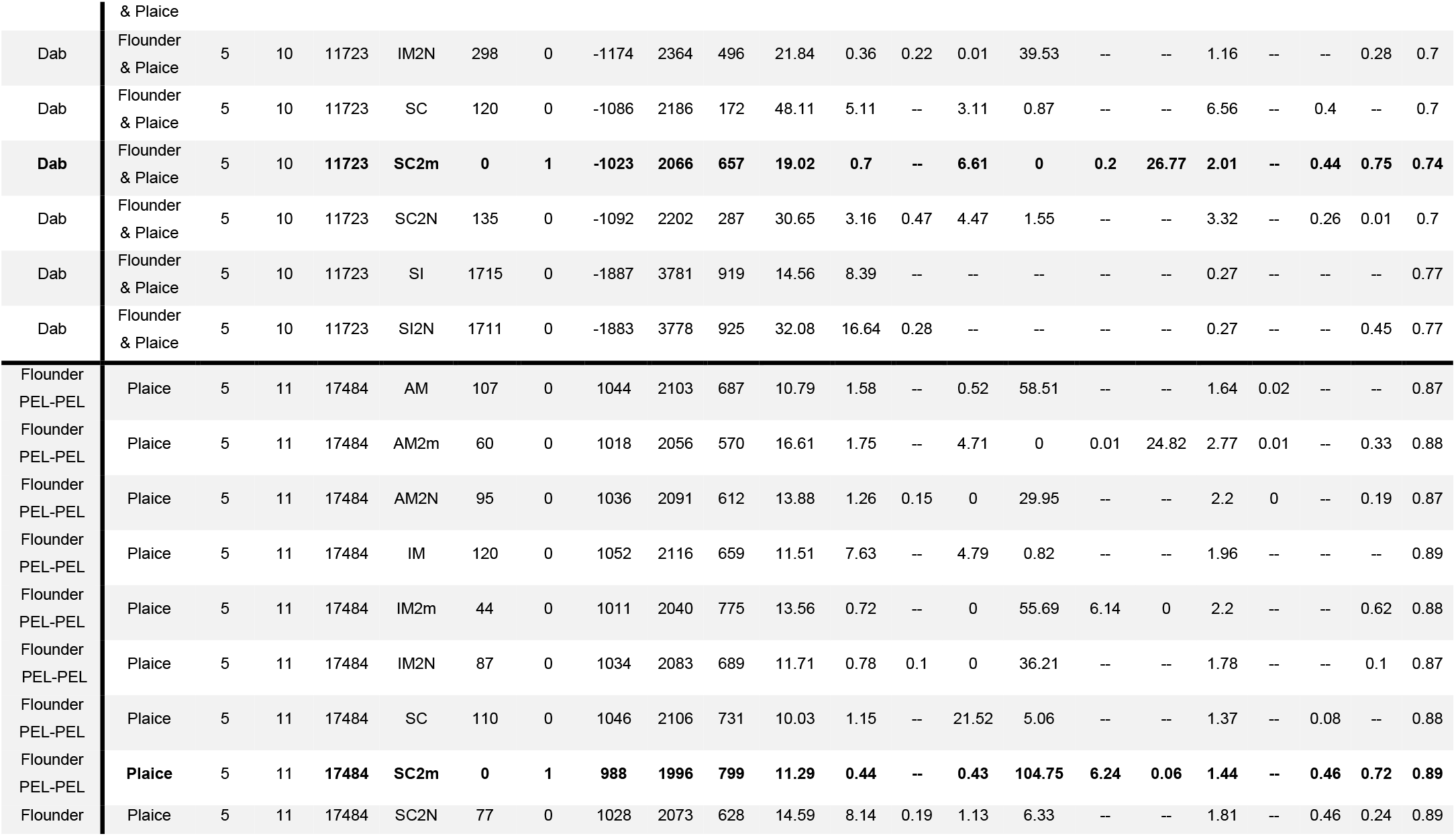
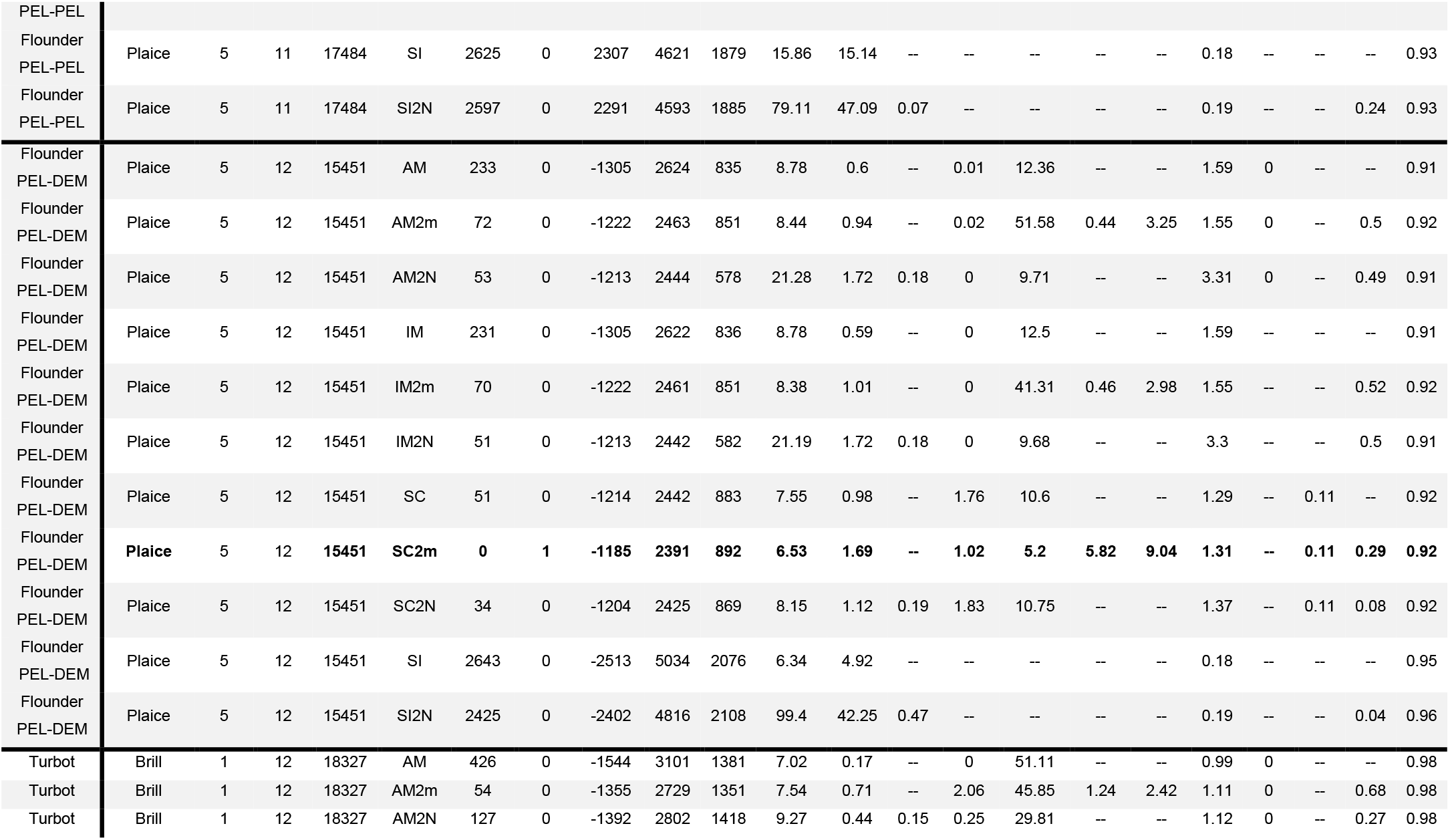
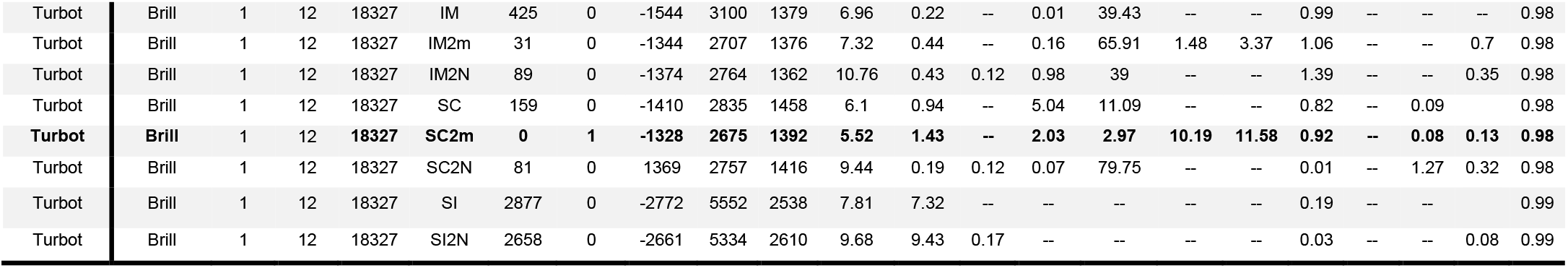
Details of the pairwise inference performed with δaδi for the best fit of each tested models. The first six columns show the focal species, the outgroup species to polarise the alleles, the pairwise sampling sites (site 1 and site 2), the number of SNPs and the different models. The next four columns provide the detailed statistics used for model selection: the differences with the best model (Δ_AIC_), the weighted AIC (more details in Rougeux et al, 2017), the likelihood (lh.) and the AIC. The remaining columns shows the parameters estimated relatively to θ: the effective population size of site 1 and site 2 respectively (N_E_1 & N_E_2), the migration rate (from site 1 to site 2 – m_1>2_ and from site 2 to site 1 – m_2>1_), the reduced migration rate (mr1>2 and mr2>1), the time of split between the populations (T_S_), time of ancestral migration (T_AM_), the time of secondary contact (T_SC_) and the proportion of loci with reduced migration rate or reduced effective size (P/Q) and the proportion of loci correctly oriented by the outgroup. The best model for each species treated in the study is highlighted in bold.

**Table S8:** Transformation of the *δaδi* parameters into meaningful biological estimates using a mutation rate of 1 x 10^-8^ and a generation time of 3.5 years in all species. In order of appearance, for each species and each model, with the transformation of *θ* into ancestral effective size (N_A_), the effective population size of the Baltic Sea and the North Sea lineage (N_E_1 and N_E_2 respectively), the migration rates (m_1>2_ and m_2>1_), the reduced migration rate (the mr_1>2_ and mr_2>1_), the time of split in kyear (T_S_), the time of ancestral migration (T_AM_) and the time of secondary contact (T_SC_)

| Species | models | $N_A$ | $N_{E1}$ | $N_{E2}$ | $m_{1>2}$ | $m_{2>1}$ | $mr_{1>2}$ | $mr_{2>1}$ | $T_S$ | $T_{AM}$ | $T_{SC}$ |
| --- | --- | --- | --- | --- | --- | --- | --- | --- | --- | --- | --- |
| Plaice | AM | 5718 | 655529 | 44491 | 0.19 | 0.00 | -- | -- | 48.80 | 0.51 | -- |
| Plaice | AM2m | 5431 | 39184 | 27780 | 0.01 | 1.30 | 0.08 | 0.01 | 54.11 | 0.00 | -- |
| Plaice | AM2N | 4572 | 74651 | 58151 | 0.00 | 0.33 | -- | -- | 63.30 | 0.25 | -- |
| Plaice | IM | 5350 | 628512 | 42191 | 0.15 | 0.01 | -- | -- | 56.08 | -- | -- |
| **Plaice** | **IM2m** | **5358** | **46218** | **23281** | **0.01** | **1.34** | **0.03** | **0.00** | **51.09** | **--** | **--** |
| Plaice | IM2N | 5340 | 57846 | 18973 | 0.00 | 0.37 | -- | -- | 60.59 | -- | -- |
| Plaice | SC | 5408 | 31342 | 15046 | 0.12 | 0.38 | -- | -- | 46.49 | 1.51 | -- |
| Plaice | SC2m | 5592 | 46686 | 41227 | 0.02 | 1.14 | 0.11 | 0.00 | 2.12 | -- | 48.07 |
| Plaice | SC2N | 5615 | 59119 | 10766 | 0.00 | 0.67 | -- | -- | 6.05 | -- | 53.50 |
| Plaice | SI | 14737 | 460979 | 460994 | -- | -- | -- | -- | 12.91 | -- | -- |
| Plaice | SI2N | 14741 | 545797 | 468506 | -- | -- | -- | -- | 13.11 | -- | -- |
| Dab | AM | 4587 | 71110 | 671 | 0.00 | 0.65 | -- | -- | 30.56 | 0.00 | -- |
| Dab | AM2m | 7320 | 88938 | 2515 | 0.00 | 0.51 | 0.04 | 0.00 | 57.81 | 0.01 | -- |
| Dab | AM2N | 4108 | 96284 | 2132 | 0.00 | 0.36 | -- | -- | 34.31 | 0.02 | -- |
| Dab | IM | 4604 | 68945 | 1025 | 0.00 | 0.43 | -- | -- | 30.71 | -- | -- |
| Dab | IM2m | 9484 | 98373 | 1697 | 0.00 | 0.78 | 0.03 | 0.00 | 55.34 | -- | -- |
| Dab | IM2N | 4234 | 92478 | 1503 | 0.00 | 0.47 | -- | -- | 34.44 | -- | -- |
| Dab | SC | 1467 | 70571 | 7501 | 0.11 | 0.03 | -- | -- | 67.36 | -- | 4.12 |
| **Dab** | **SC2m** | **5604** | **106594** | **3913** | **0.06** | **0.00** | **0.00** | **0.02** | **78.89** | **--** | **17.28** |
| Dab | SC2N | 2448 | 74905 | 7726 | 0.09 | 0.03 | -- | -- | 56.85 | -- | 4.40 |
| Dab | SI | 7836 | 114067 | 65778 | -- | -- | -- | -- | 14.74 | -- | -- |
| Dab | SI2N | 7890 | 253122 | 131292 | -- | -- | -- | -- | 14.70 | -- | -- |
| Flounder PEL-PEL | AM | 3931 | 42417 | 6205 | 0.01 | 0.74 | -- | -- | 45.18 | 0.60 | -- |
| Flounder PEL-PEL | AM2m | 3259 | 54134 | 5704 | 0.07 | 0.00 | 0.00 | 0.04 | 63.16 | 0.12 | -- |
| Flounder PEL-PEL | AM2N | 3498 | 48543 | 4418 | 0.00 | 0.43 | -- | -- | 53.83 | 0.05 | -- |
| Flounder PEL-PEL | IM | 3768 | 43333 | 28742 | 0.06 | 0.01 | -- | -- | 51.82 | -- | -- |
| Flounder PEL-PEL | IM2m | 4431 | 60090 | 3178 | 0.00 | 0.63 | 0.07 | 0.00 | 68.36 | -- | -- |
| Flounder PEL-PEL | IM2N | 3938 | 46109 | 3065 | 0.00 | 0.46 | -- | -- | 48.98 | -- | -- |
| Flounder PEL-PEL | SC | 4182 | 41925 | 4797 | 0.26 | 0.06 | -- | -- | 40.17 | -- | 2.38 |
| **Flounder PEL-PEL** | **SC2m** | **4571** | **51613** | **2027** | **0.00** | **1.15** | **0.07** | **0.00** | **46.02** | **--** | **14.75** |
| Flounder PEL-PEL | SC2N | 3594 | 52453 | 29257 | 0.02 | 0.09 | -- | -- | 45.55 | -- | 11.46 |
| Flounder PEL-PEL | SI | 10748 | 170435 | 162253 | -- | -- | -- | -- | 13.63 | -- | -- |
| Flounder PEL-PEL | SI2N | 10783 | 853041 | 507773 | -- | -- | -- | -- | 14.22 | -- | -- |
| Flounder PEL-DEM | AM | 5407 | 47452 | 3240 | 0.00 | 0.11 | -- | -- | 60.33 | 0.00 | -- |
| Flounder PEL-DEM | AM2m | 5509 | 46485 | 5167 | 0.00 | 0.47 | 0.00 | 0.00 | 59.91 | 0.01 | -- |
| Flounder PEL-DEM | AM2N | 3744 | 79651 | 6451 | 0.00 | 0.13 | -- | -- | 86.87 | 0.00 | -- |
| Flounder PEL-DEM | IM | 5409 | 47474 | 3205 | 0.00 | 0.12 | -- | -- | 60.32 | -- | -- |
| Flounder PEL-DEM | IM2m | 5508 | 46169 | 5556 | 0.00 | 0.37 | 0.00 | 0.00 | 59.78 | -- | -- |
| Flounder PEL-DEM | IM2N | 3764 | 79757 | 6489 | 0.00 | 0.13 | -- | -- | 86.88 | -- | -- |
| Flounder PEL-DEM | SC | 5712 | 43138 | 5601 | 0.02 | 0.09 | -- | -- | 51.75 | -- | 4.60 |
| **Flounder PEL-DEM** | **SC2m** | **5774** | **37732** | **9766** | **0.01** | **0.05** | **0.05** | **0.01** | **52.92** | **--** | **4.38** |
| Flounder PEL-DEM | SC2N | 5622 | 45837 | 6318 | 0.02 | 0.10 | -- | -- | 54.02 | -- | 4.47 |
| Flounder PEL-DEM | SI | 13439 | 85176 | 66076 | -- | -- | -- | -- | 16.66 | -- | -- |
| Flounder PEL-DEM | SI2N | 13643 | 1356145 | 576356 | -- | -- | -- | -- | 17.98 | -- | -- |
| Turbot | AM | 7537 | 52889 | 1272 | 0.00 | 0.34 | -- | -- | 52.15 | 0.00 | -- |
| Turbot | AM2m | 7371 | 55568 | 5209 | 0.01 | 0.31 | 0.01 | 0.00 | 57.27 | 0.02 | -- |
| Turbot | AM2N | 7736 | 71692 | 3437 | 0.00 | 0.19 | -- | -- | 60.53 | 0.00 | -- |
| Turbot | IM | 7522 | 52376 | 1654 | 0.00 | 0.26 | -- | -- | 52.21 | -- | -- |
| Turbot | IM2m | 7510 | 54970 | 3268 | 0.00 | 0.44 | 0.01 | 0.00 | 55.95 | -- | -- |
| Turbot | IM2N | 7434 | 79975 | 3213 | 0.01 | 0.26 | -- | -- | 72.43 | -- | -- |
| Turbot | SC | 7957 | 48566 | 7473 | 0.03 | 0.07 | -- | -- | 45.85 | -- | 4.86 |
| **Turbot** | **SC2m** | **7596** | **41962** | **10889** | **0.07** | **0.08** | **0.01** | **0.02** | **49.17** | **--** | **4.07** |
| Turbot | SC2N | 7724 | 72924 | 1456 | 0.00 | 0.52 | -- | -- | 0.67 | -- | 68.55 |
| Turbot | SI | 13847 | 108104 | 101335 | -- | -- | -- | -- | 18.25 | -- | -- |
| Turbot | SI2N | 14239 | 137817 | 134242 | -- | -- | -- | -- | 3.33 | -- | -- |

**Figure S7:**
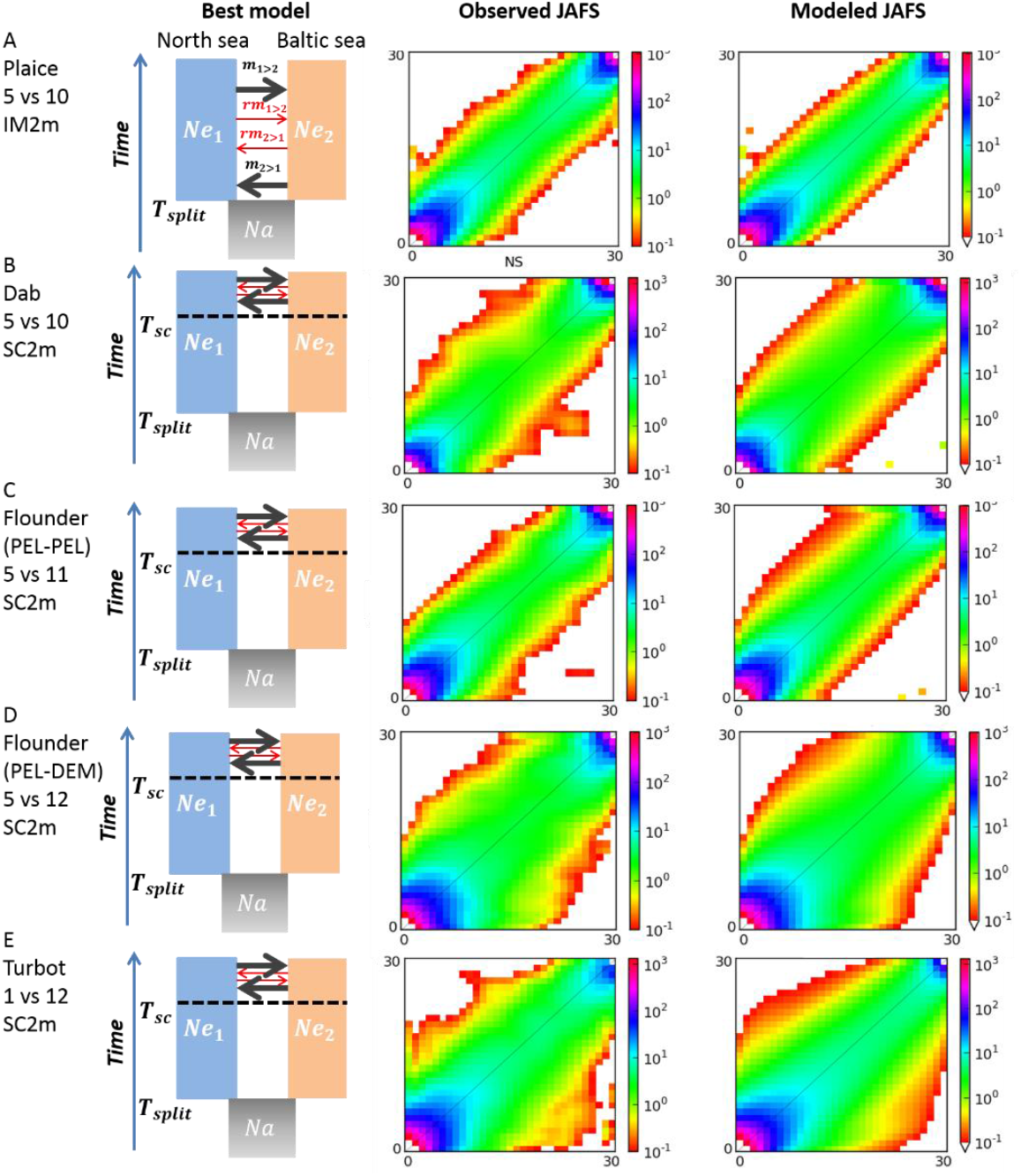
Results from the best model fit for the North Sea (y axis of the JAFS) and Baltic Sea (x axis of the JAFS) pairwise inference for A) the plaice, B) the dab, C) the pelagic flounder D) the pelagic vs demersal flounder and D) the turbot. Left: representation of best model based on wAIC (SC2m for all species expect the plaice with IM2m). Middle: the observed joint allele frequency spectrum showing the number of SNPs (color scale on the right of the spectrum) per pixel of derived allele count. Right: The maximum-likelihood JAFS obtained under the best model. Inference for plaice was based on keeping information for only one SNP within each putative structural variant.

### Tests for selection

**Figure S8:**
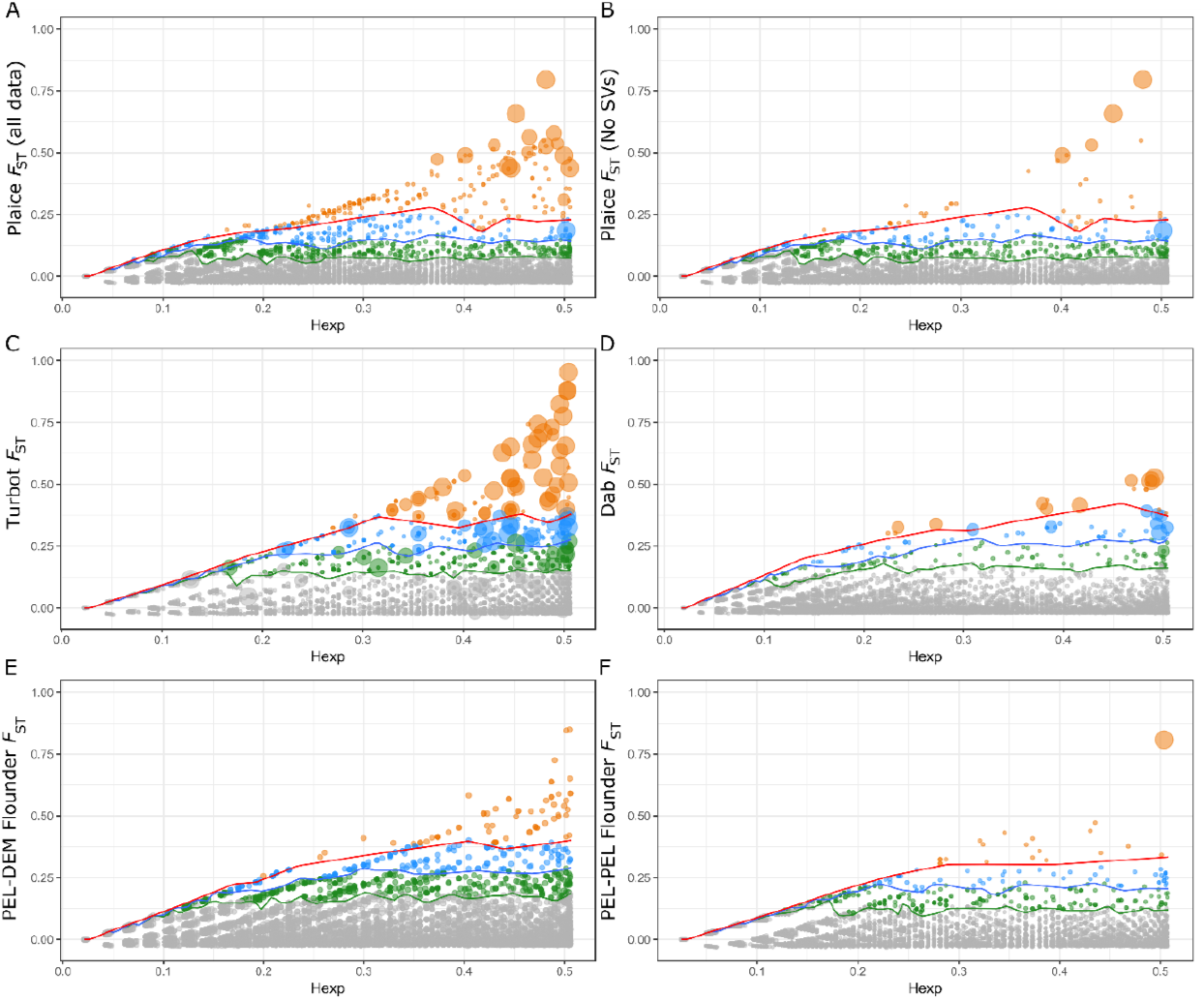
Fst genome scans. The straight lines correspond to the envelope of neutral divergence for the 5%, 1% and 0.1% upper quantile of differentiation (in green, blue and red, respectively). The colours of the dots correspond to the genome scan outlier (green = top 5%, blue = top 1% and orange = top 0.1%) and the largest dots correspond to the environmental outliers for A-B) plaice, respectively with and without the SVs, C) turbot, D) dab, E) the pelagic and demersal flounder, and F) the pelagic flounder only.

**Figure S9:**
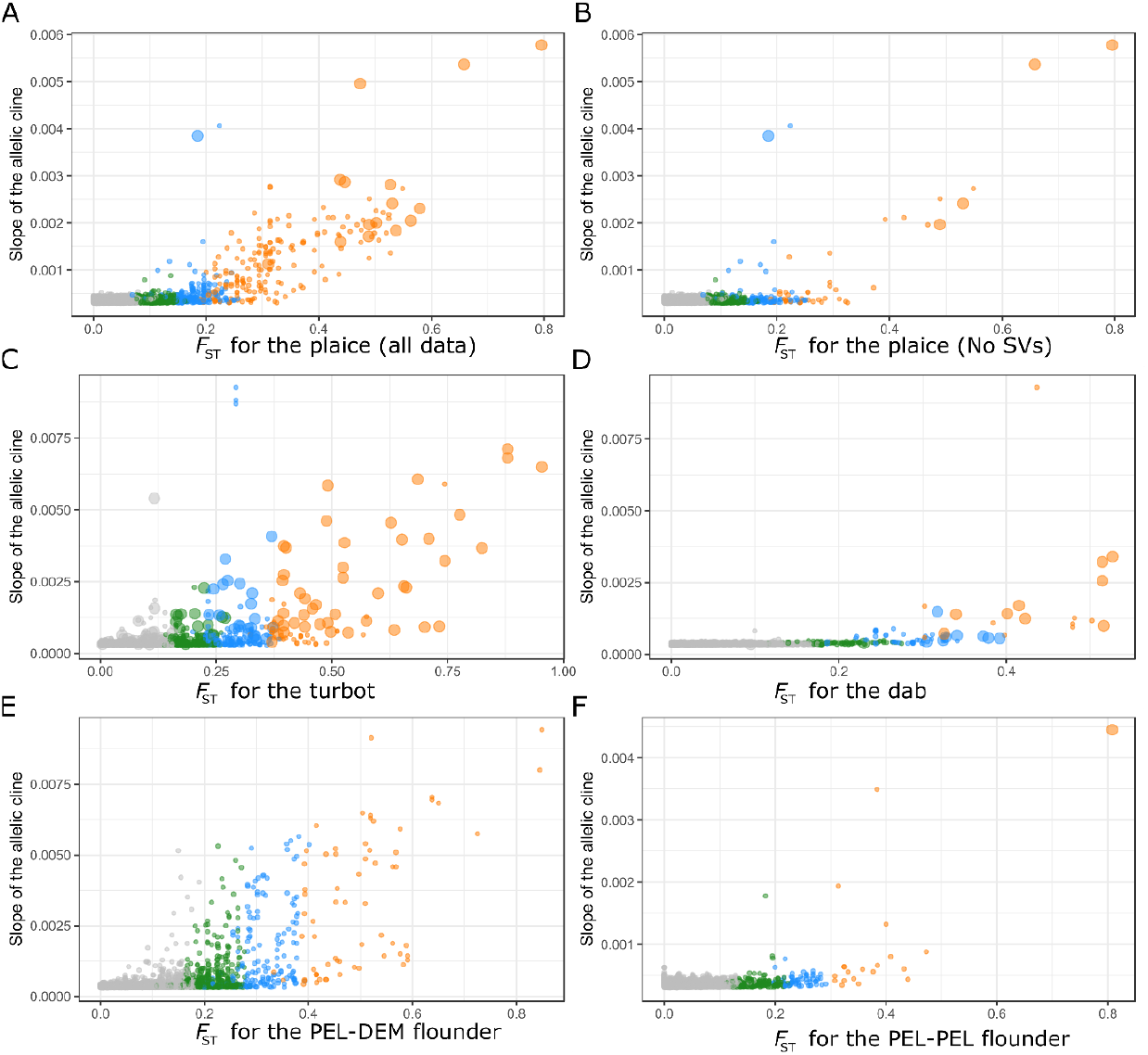
Correlation between the *F*_ST_ and the estimates of allelic cline slope. The colours correspond to the genome scan outliers (green = top 5%, blue = top 1% and orange = top 0.1%) and the largest dots correspond to the environmental outliers for A-B) plaice with and without the SVs, respectively, C) turbot, D) dab E) the pelagic and demersal flounder, and F) the pelagic flounder only.

**Figure S10:**
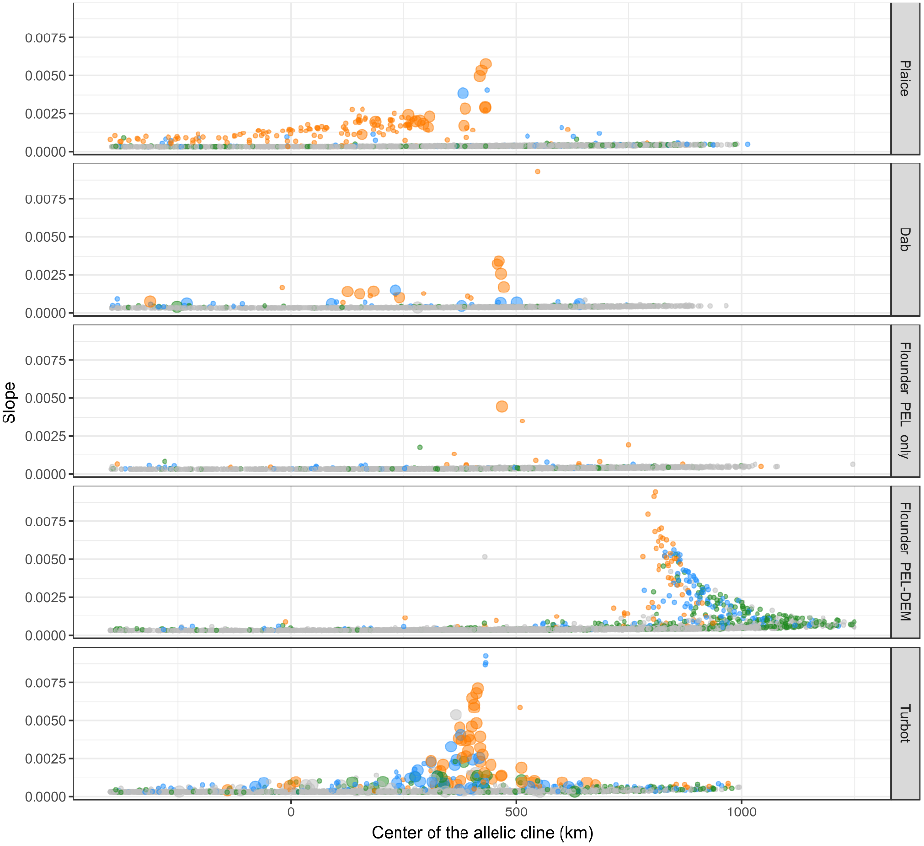
Geography of clines based the slope of allele frequencies for individual markers as a function of the distance of the cline center from the North Sea for each species. The colours correspond to the genome scan outliers (green = top 5%, blue = top 1% and orange = top 0.1%) and the largest dots correspond to the environmental outliers

**Figure S11:**
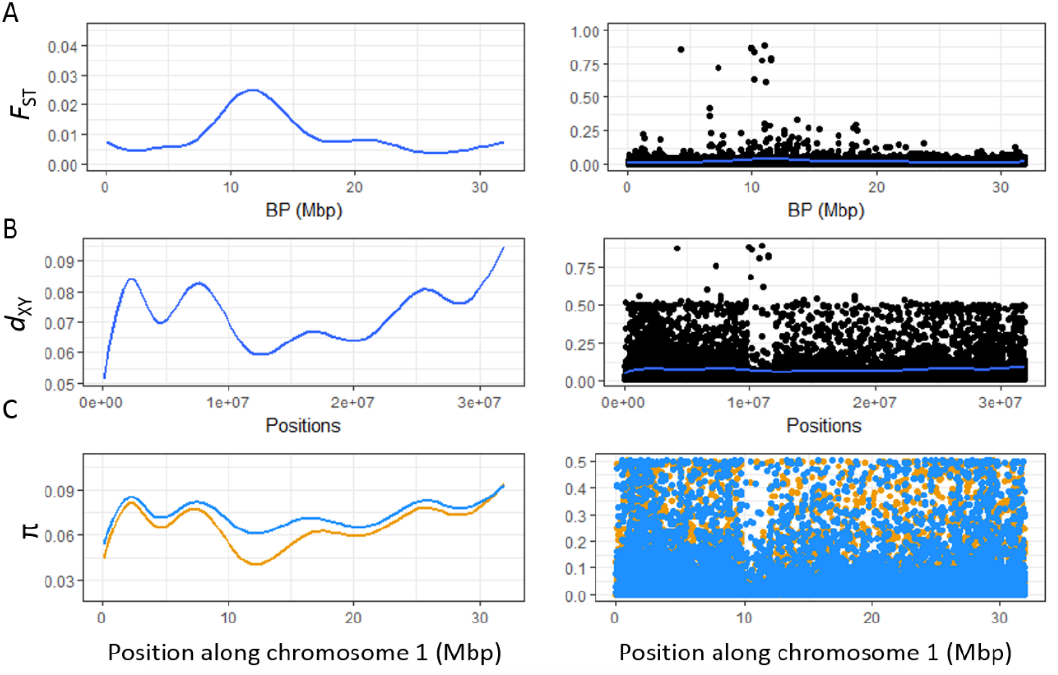
Summary statistics of the turbot North Sea – Baltic Sea divergence (*F*_ST_ and *d*_XY_) and genetic diversity (π) for the North Sea (blue) and the Baltic Sea (yellow) along chromosome 1. The graphs on the left show the smoothed average statistics over bins of 100kb and the graphs on the right show the individual SNP estimates. The peak of F_ST_ localized in the center of the chromosome co-localized with a drop in d_XY_ and diversity in both populations corresponding roughly to the location of the centromere.

## Material and Methods

### Sampling

**Table S9:** Sampling details. The sampling locations are shown together with sampling site numbers used in figures. Site 1 corresponds to the most western site, and site 12 to the most north-eastern site within the Baltic Sea. For each site, number of sequenced individuals (‘Sample size’), month and year of sampling are shown. The last column refers to studies where the same samples were analysed with different molecular approaches.

| Species | area | Site | Sample size | Year | Publication |
| --- | --- | --- | --- | --- | --- |
| Dab | North Sea | 5 | 31 | Feb. 2016 |  |
|  | Kattegat | 6 | 25 | Feb. 2016 |  |
|  | Great belt | 7 | 25 | Feb. 2016 |  |
|  | Øresund | 8 | 25 | Mar. 2016 |  |
|  | South-West of Baltic | 9 | 25 | Apr. 2016 |  |
|  | Bornholm | 10 | 17 | Mar. 2017 |  |
| Plaice | North Sea | 5 | 30 | Feb. 2016 |  |
|  | Skagerrak | 6 | 25 | Feb. 2017 |  |
|  | Kattegat | 6 | 25 | Feb. 2017 |  |
|  | Great belt | 7 | 25 | Feb. 2016 |  |
|  | Øresund | 8 | 13 | Mar. 2016 |  |
|  |  |  | 12 | Mar. 2017 |  |
|  | South-West of Baltic | 9 | 25 | Apr. 2016 |  |
|  | Bornholm | 10 | 25 | Mar. 2017 |  |
| Flounder | North Sea | 5 | 30 | Feb. 2016 |  |
|  | Kattegat | 6 | 25 | Feb. 2017 |  |
|  | Great belt | 7 | 25 | Feb. 2016 |  |
|  | Øresund | 8 | 25 | Mar. 2016 |  |
|  | South-West of Baltic | 9 | 25 | Apr. 2016 |  |
|  | Bornholm | 10 | 25 | Mar. 2016 |  |
|  | South-East of Baltic | 11 | 25 | Mar. 2016 |  |
|  | North of Baltic | 12 | 14 | June 2016 |  |
|  |  |  | 20 | June 2003 | Hemmer-Hansen *et al.* (2007) |
| Turbot | English Channel | 1 | 25 | Dec. 2009 | Vandamme *et al.* (2014) |
|  | South-West North Sea | 2 | 25 | Nov. 2009 | Vandamme *et al.* (2014) |
|  | North-West North Sea | 3 | 11 | Nov. 2010 | Vandamme *et al.* (2014) |
|  | South-East North Sea | 4 | 26 | Nov. 2010 | Vandamme *et al.* (2014) |
|  | North East of North Sea | 5 | 19 | Feb. 2016 |  |
|  |  |  | 6 | Oct. 2010 | Vandamme *et al.* (2014) |
|  | Kattegat | 6 | 25 | Feb. 2017 |  |
|  | Great belt | 7 | 25 | Feb. 2016 |  |
|  | Øresund | 8 | 11 | Mar. 2016 |  |
|  |  |  | 14 | Mar. 2013 | Vandamme *et al.* (2014) |
|  | South-West of Baltic | 9 | 25 | Apr. 2016 |  |
|  | Bornholm | 10 | 27 | Mar. 2016 |  |
|  | South-East of Baltic | 11 | 14 | Mar. 2016 |  |
|  | North of Baltic | 12 | 2 | June 2016 |  |
|  |  |  | 20 | Aug. 1999 | Nielsen *et al.* (2003) |
| Brill | North Sea | 5 | 8 | Feb. 2016 |  |

### Bioinformatics

**Figure S12:**
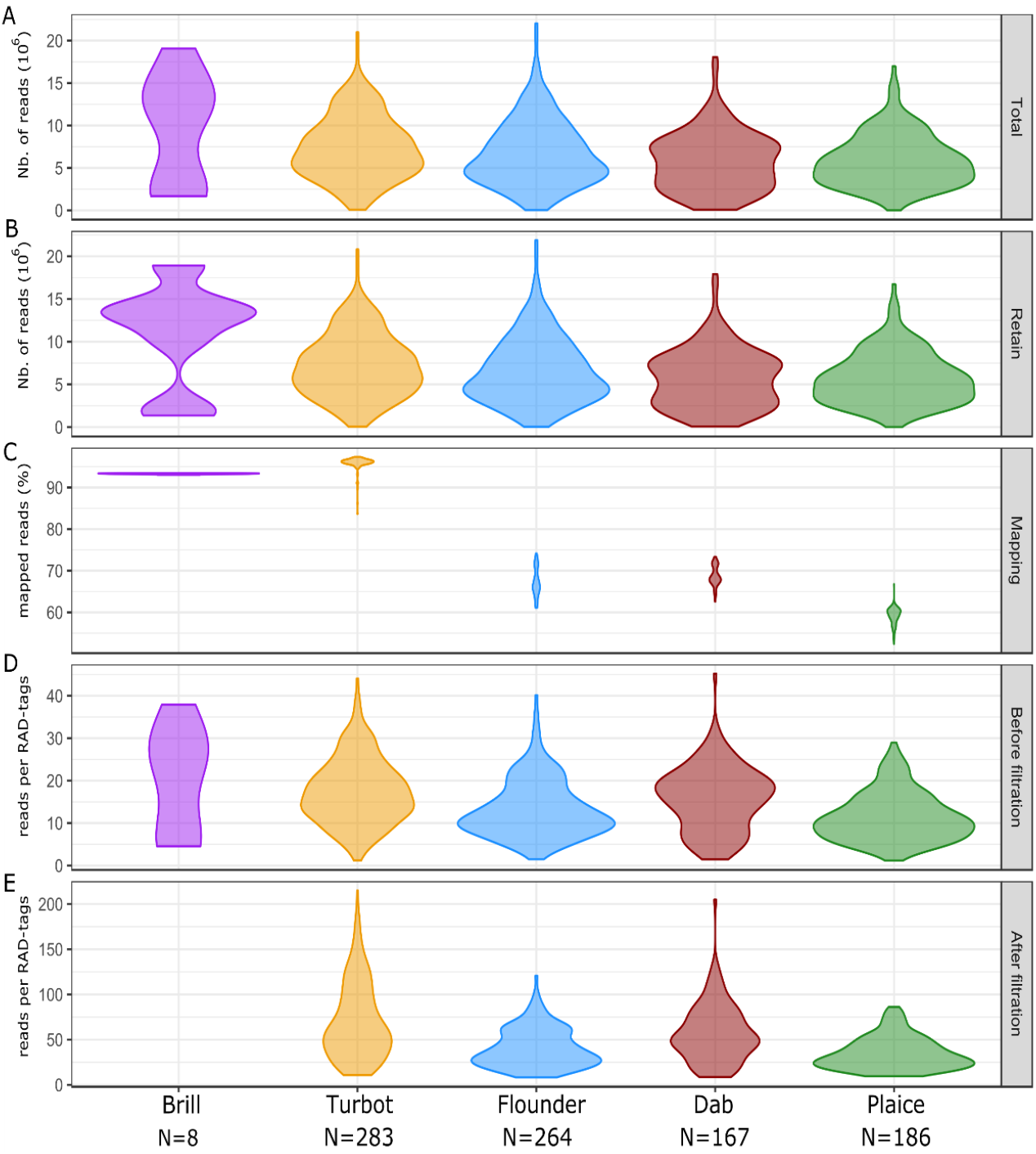
Violin plots showing the distribution of individual quality statistics for each studied species with A) the total number of reads after sequencing, B) the number of reads with phred33 quality scores above 10, c) the percentage of reads mapped back to the reference genomes, D) the average coverage per RAD-tag and E) the average coverage per RAD-tag after filtration.

**Figure S12a:**
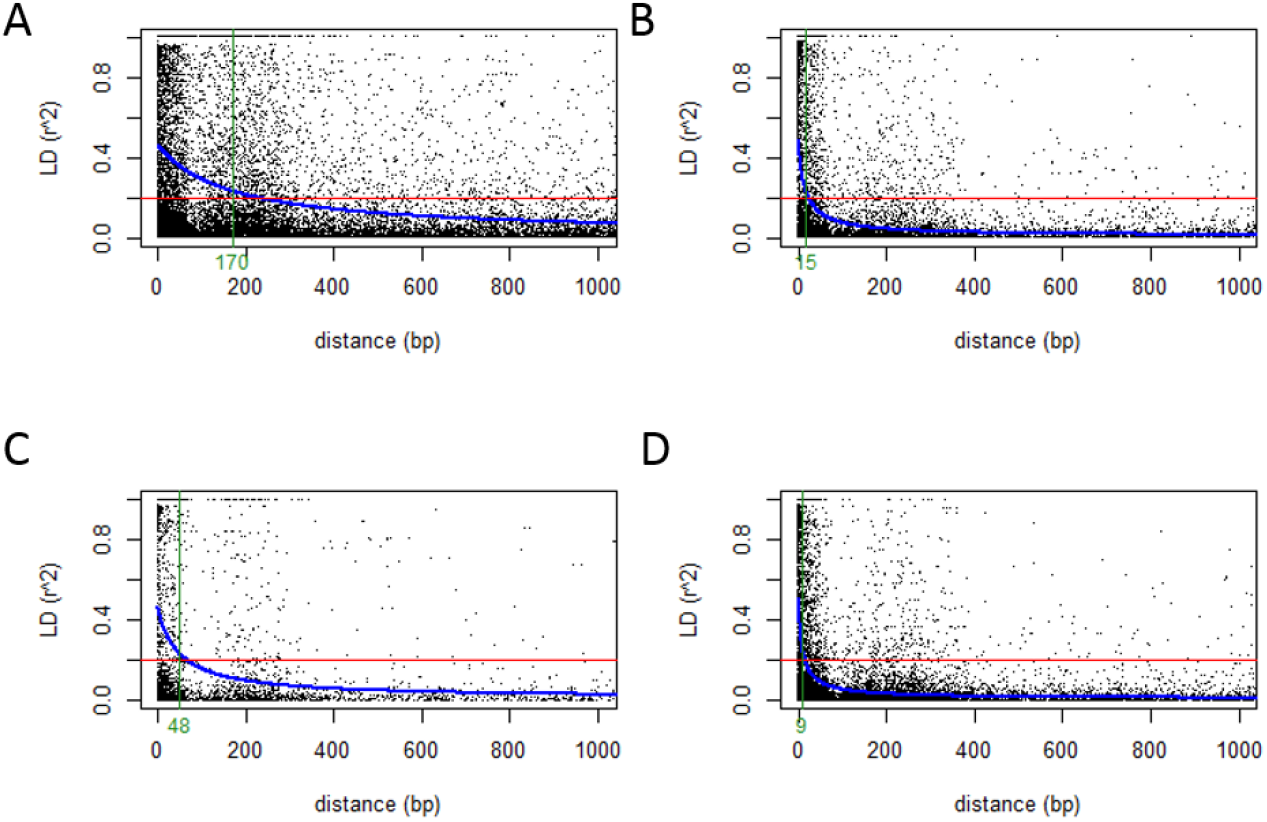
LD decay inference from the ddRAD-data for A) turbot, B) flounder C) dab and D) plaice. The red line shows the LD value after which SNPs can be considered as independent and the green line shows distance after which more than 95% of the comparisons have a LD value bellow 0.2.

